# A novel G protein-biased and subtype selective agonist for a G protein-coupled receptor discovered from screening herbal extracts

**DOI:** 10.1101/2019.12.22.883686

**Authors:** Bingjie Zhang, Simeng Zhao, Dehua Yang, Yiran Wu, Ye Xin, Haijie Cao, Xi-Ping Huang, Xiaoqing Cai, Wen Sun, Na Ye, Yueming Xu, Yao Peng, Suwen Zhao, Zhi-Jie Liu, Guisheng Zhong, Ming-Wei Wang, Wenqing Shui

**Author notes:** To whom correspondence should be addressed to: Wenqing Shui Ming-Wei Wang Guisheng Zhong. Equal contribution (Bingjie Zhang, Simeng Zhao).

## Abstract

Subtype selectivity and functional bias are vital in current drug discovery for G protein-coupled receptors (GPCRs) as selective and biased ligands are expected to yield drug leads with optimal on-target benefits and minimal side-effects. However, structure-based design and medicinal chemistry exploration remain challenging in part because of highly conserved binding pockets within subfamilies. Herein, we present an affinity mass spectrometry approach for screening herbal extracts to identify active ligands of a GPCR, the 5-HT_2C_ receptor. Using this method, we discovered a naturally occurring aporphine 1857 that displayed strong selectivity for activating 5-HT_2C_ without activating the 5-HT_2A_ or 5-HT_2B_ receptors. Remarkably, this novel ligand exhibited exclusive bias towards G protein signaling for which key residues were identified, and it showed comparable *in vivo* efficacy for food intake suppression and weight loss as the anti-obesity drug, lorcaserin. Our study establishes an efficient approach to discovering novel GPCR ligands by exploring the largely untapped chemical space of natural products.

## Introduction

Belonging to the superfamily of G protein-coupled receptors (GPCRs), the serotonin (5-hydroxytryptamine, 5-HT) receptors mediate a plethora of physiological processes in the brain and the periphery^1^. The human genome encodes thirteen 5-HT receptors that exert the biological effects of serotonin and eight are drug targets for the treatment of obesity, migraine, anxiety, depression and hypertension^1, 2^. Among them, the serotonin 2C receptor (5-HT_2C_) is recognized as a promising therapeutic target for obesity and central nervous system (CNS) disorders, such as epilepsy, schizophrenia and drug abuse^2–4^. The value of 5-HT_2C_ in anti-obesity medication development is manifested by the FDA-approved drug lorcaserin, a 5-HT_2C_ selective agonist^1^. Moreover, the efficacy of lorcaserin in treatment of nicotine addiction is currently being evaluated clinically^5^.

The development of 5-HT_2C_ agonists as potential anti-obesity and anti-psychotic medications requires high selectivity over other subfamily members, the 5-HT_2A_ and 5-HT_2B_ receptors, whose activation is associated with hallucination^6^ and cardiac valvulopathy^7, 8^. For example, due to their off-target activities at 5-HT_2B_, the non-selective serotoninergic drugs, fenfluramine and pergolide, were withdrawn from markets, and the drug cabergoline has been restricted^8–10^. Even the safety of lorcaserin has been questioned due to its moderate selectivity (∼100-fold) over 5-HT_2B_^11, 12^. However, developing subtype selective agonists for 5-HT_2C_ is challenging owing to the highly conserved ligand-binding pockets among the three 5-HT_2_ members^13–15^. To date, only a handful of scaffolds have been disclosed as selective 5-HT_2C_ agonists, all of which were obtained through extensive medicinal chemistry exploration^16–20^.

The concept of signaling bias or functional selectivity has recently reshaped our understanding of GPCR signaling and shifted the paradigm for GPCR drug discovery^21, 22^. Signaling bias refers to a process whereby GPCR ligands can either activate G proteins or recruit β-arrestins to mediate specific downstream signaling pathways for a given receptor^23, 24^. Biased GPCR ligands, which can trigger a specific pathway responsible for a given therapeutic effect while not activating other pathways that are implicated in side-effects, possess significant potential to become drug leads with optimal on-target benefits^25, 26^. For example, G protein-biased μ-opioid receptor agonists are potentially analgesic but have reduced side-effects (*e.g.* respiratory depression and constipation)^27, 28^. Although an increasing number of biased ligands have been discovered for different GPCRs^26, 29–32^, very few for 5-HT_2C_ have been reported. Recently, a class of compounds based on the (2-phenylcyclopropyl)methylamine scaffold synthesized by Chen *et al.* and Zhang *et al.* exhibited functional selectivity at 5-HT_2C_ with preference to G_q_-mediated calcium flux^16, 19^. However, synthesizing compounds with both signaling bias and subtype selectivity remains a major obstacle for medicinal chemists.

A rich resource for generating tool compounds and drug leads are the natural herbs, as their chemical constituents typically possess molecular architectures and bioactivities that are distinct from synthetic molecules^33, 34^. To expedite ligand discovery for various protein targets from natural products, a number of approaches have been developed, ranging from cell-based activity or biosensor-based binding assays to *in silico* screening^33, 35^. Unlike most screening platforms that examine individual pure compounds from a library, affinity mass spectrometry (MS) can directly capture and detect putative ligands from crude natural product extracts towards a protein target^36–40^. Although affinity MS has shown great potential in discovering inhibitors or modulators of enzymes and other soluble protein targets^38, 39, 41–43^, it has never been explored in GPCR ligand screening from natural products.

In this study, we adapted the affinity MS technique to discover new ligands for 5-HT_2C_ from a collection of natural product extracts. Emerging from this screen was a unique family of aporphine alkaloids, a rarely investigated chemotype for this target. Guided by the affinity MS screening data, we were able to isolate two novel aporphine ligands for pharmacological characterization. For one ligand 1857 that acted as a selective 5-HT_2C_ agonist with exclusive G protein signaling bias, key residues for 5-HT_2C_ activation were then identified by molecular docking and mutagenesis. Finally, we compared this herb-derived novel agonist against the approved drug lorcaserin for *in vivo* anti-obesity effects.

## Results

### Selection of natural herbs for affinity MS screening

To select natural herbs for affinity MS screen of 5-HT_2C_ agonists, we first assayed bioactivities of crude extracts from 15 different plants using calcium flux assay that separately measures G_q_-coupled activities of 5-HT_2C_ receptor and two close family members, 5-HT_2A_ and 5-HT_2B_ receptors. The total extracts from eight herbs showed agonist property at 5-HT_2C_ with potency spanning from nM to µM (assuming an average molecular weight of 500 Da for small molecule constituents) (Fig. 1A). Given that major constituents typically account for less than 1% of the total weight, we speculated that agonists with low nanomolar potency might be present in the extracts of *Aristolochia debilis* (AD) and *Tetradium ruticarpum* (TR) showing the highest potency at 5-HT_2C_. Moreover, among the eight herbs with 5-HT_2C_ agonism, five also activated 5-HT_2A_ and 5-HT_2B_ with similar potency (Figs. 1B-D). However, this does not exclude the possibility that individual components in the extracts may possess subtype selectivity. Therefore, these five extracts were selected for additional screening with the affinity MS approach. We first prepared the apo 5-HT_2C_ protein fused with a stabilizing partner^15^ without any mutation and staying in a homogenous monomeric conformation after purification (Supplementary Fig. 1). The purified receptor immobilized on magnetic beads through an epitope tag was then incubated with a defined compound mixture. Ligand-bound 5-HT_2C_ complexes were enriched by magnetic separation from the solution phase. Bound ligands were dissociated from the protein target and subjected to liquid chromatography coupled to high-resolution mass spectrometry (LC-HRMS) analysis (Fig. 1E). Another purified GPCR (hydroxyl carboxylic acid receptor 2, HCA2) was immobilized and processed in the same manner to serve as a negative control (Supplementary Fig. 1). Quantitative comparison of the extracted ion chromatograms of individual compounds detected in the target versus the control allowed us to distinguish specific 5-HT_2C_ ligands from non-specific binders (Fig. 1E). This method was first validated using a mixture of known 5-HT_2C_ ligands and unrelated compounds. All agonists and antagonists with high affinity (K_i_ <50 nM) were associated with the receptor in the affinity MS assay, while none of the unrelated compounds were qualified as hits (Supplementary Fig. 2).

**Figure 1.**
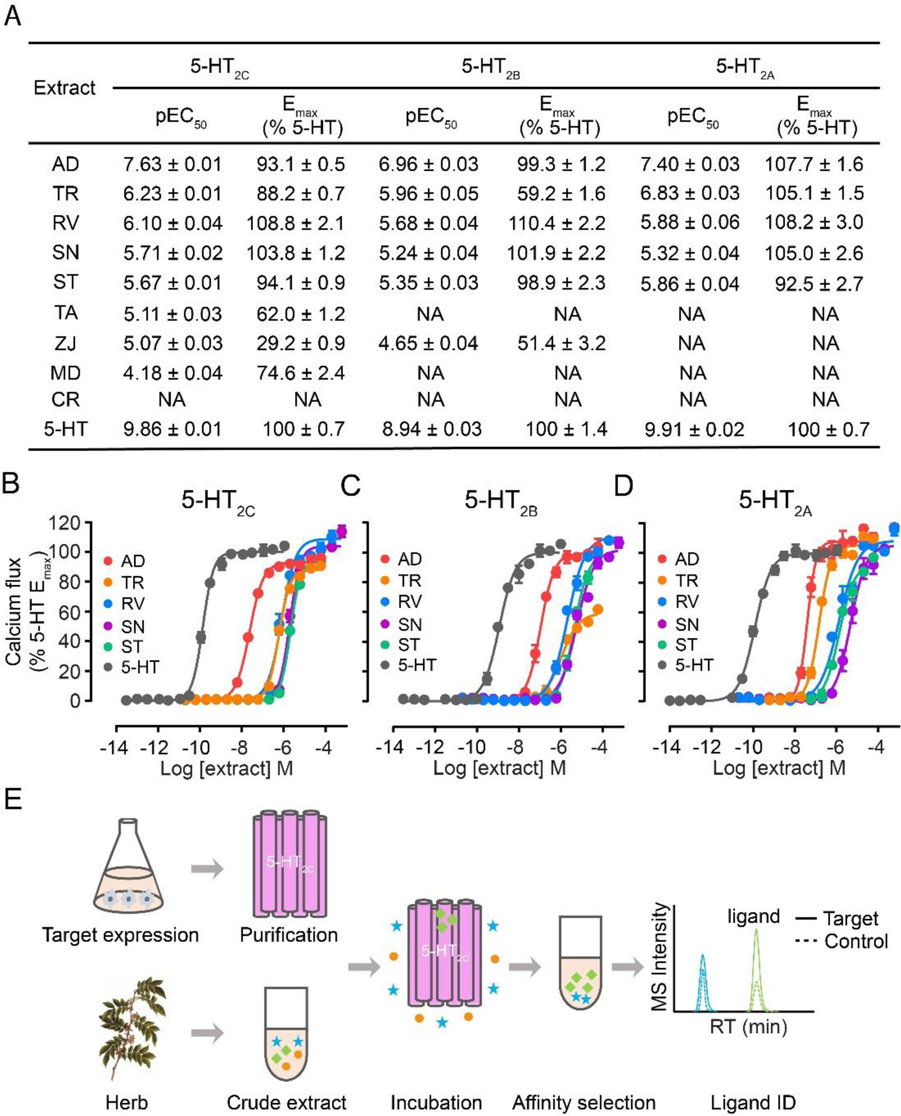
Herbal extracts showing G_q_-mediated activity at 5-HT_2C_ were selected for affinity MS screening. (A) Calcium mobilization elicited by three 5-HT_2_ subtypes treated with different herbal extracts or 5-HT. Full names of the herbs used are listed in Methods. NA, not active. (B-D) Dose-response characteristics of calcium mobilization elicited by top 5 extracts or 5-HT at 5-HT_2C_, 5-HT_2B_ and 5-HT_2A_. Crude extract concentrations were calculated from weight, assuming an average molecular weight of 500 Da for small molecule constituents. (E) The workflow of affinity MS-based 5-HT_2C_ ligand screening. A putative ligand (green) and a non-specific binder (blue) are distinguished based on the MS intensity of each compound detected in the 5-HT_2C_ target *vs.* control.

### Identification of new 5-HT_2C_ ligands from herbal extracts

The established affinity MS workflow was first applied to screening 5-HT_2C_ ligands from crude extracts of AD and TR that displayed the highest potency among all natural herbs assayed. The purified receptor was incubated with either extract and underwent the same affinity MS procedure as described above. A targeted metabolomics data mining strategy previously developed by us^37^ was implemented to process the affinity MS screening data for individual extracts. Screening hits were selected if their mean binding indexes (BI) were above 2.0 (*P* < 0.05, *n* = 4)^44, 45^, and their chemical structures were assigned by matching the LC-MS features with compounds registered in a natural herb database (TCMHD)^46^. Unexpectedly, serotonin, the natural ligand for 5-HT receptors, and its analogue 5-methoxy-*N*, *N*-dimethyltryptamine (5-MeO-DMT) known as 5-HT_2C_ agonist^47^ turned out to be top-ranking hits from screening the two extracts. (Supplementary Fig. 3A-3C, Supplementary Table 1, Supplementary Note 1). Furthermore, serotonin alone or serotonin combined with 5-MeO-DMT accounted for 100% of the overall 5-HT_2C_ activity of AD and TR (Supplementary Fig. 3D-3E, Supplementary Fig. 4). As serotonin was also detected in RV and SR extracts, these four herbs were abandoned for further experimentation (Supplementary Note 1).

Extract of the fifth candidate herb *Stephania tetrandra* (ST) with appreciable 5-HT_2C_ agonism (EC_50_ = 2.13 µM) was subjected to the affinity MS screen for putative ligands (Fig. 2A) which gave rise to 12 initial hits (Fig. 2B, Supplementary Table 2). To increase the chance of capturing ligands of low abundances in the original extract, we fractionated the extract and screened each fraction separately (Supplementary Fig. 5). As expected, screening three fractions altogether allowed us to interrogate a lot more constituents (1364 assigned features in total) and identify more putative ligands than screening the crude extract alone (Fig. 2C). A close inspection of the assigned structures for all putative ligands from different screens revealed a cluster of aporphine alkaloids that possess a characteristic tetracyclic framework (Fig. 2D, Supplementary Table 3). Compounds in this alkaloid subclass have been rarely associated with 5-HT_2C_^48^. Among the eight identified aporphine ligands, only nuciferine (compound 15856) was reported to be a 5-HT_2C_ antagonist^49^ while the rest have not been associated with 5-HT_2C_.

**Figure 2.**
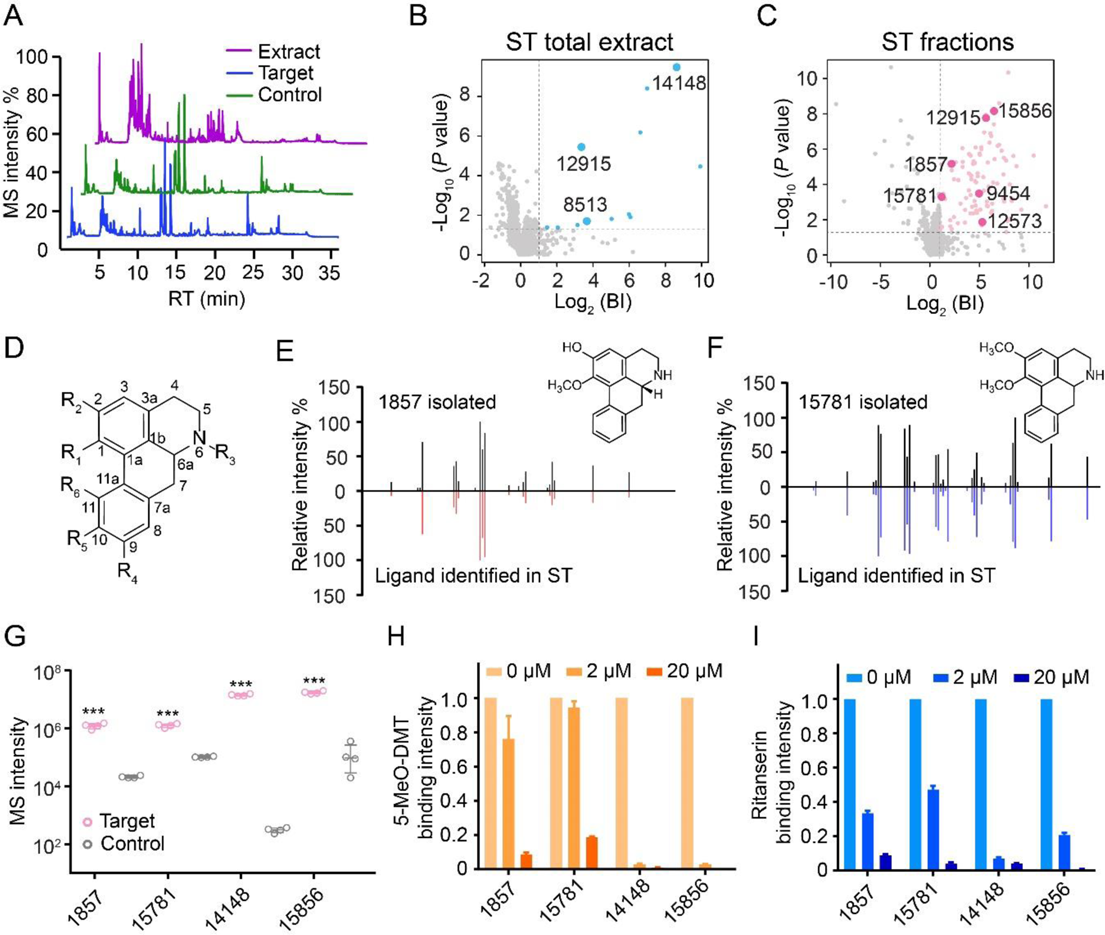
Identification of aporphines active at 5-HT_2C_ from crude and fractionated extracts of *Stephania tetrandra* (ST). (A) Representative LC-MS chromatograms of ST crude extract, 5-HT_2C_ target and control. (B,C) Initial hits from screening ST crude extract (blue dots) or ST extract fractions (pink dots) by affinity MS combined with metabolomics. Aporphines are annotated with larger dots. BI, binding index. (D) Scaffold of the aporphines identified in this study (specific structures listed in Table S3). (E,F) Structural validation of 1857 and 15781 by MSMS analysis. (G) Validation of ligand binding to purified 5-HT_2C_ by pure compounds using affinity MS binding assay. The MS intensity of each ligand was significantly higher in 5-HT_2C_ than that of control. (\*\*\**P* <0.001, n=4). (H,I) Competition of 5-MeO-DMT or ritanserin binding to purified 5-HT_2C_ with increasing concentrations of each aporphine. MS intensity of 5-MeO-DMT or ritanserin bound to purified 5-HT_2C_ was normalized to that in the absence of any aporphine. Data were obtained from two independent experiments in technical duplicate. Error bars represent SEM.

To verify their binding ability and bioactivity, we purchased pure standards for two ligands (14148 and 15856) while others are not commercially available. We then tried to isolate them directly from the extracts. Guided by the initial affinity MS screening data which pinpointed the expected retention time and accurate mass of the compounds of interest (Supplementary Table 3), we were able to isolate two new putative ligands 1857 and 15781 (Supplementary Fig. 6) and their chemical structures were confirmed by 1D and 2D NMR analysis (Supplementary Note 2). Optical rotation measurement indicated 20 that 1857 ([α_D_^20^] = −92°) is an *R* stereoisomer whereas 15781 ([α_D_^20^] = −1.4°) is a racemic mixture. When we overlaid the MSMS spectra of the isolated compound and the putative ligand identified in the affinity MS screen, almost identical MSMS fragmentation patterns strongly corroborated the structural identity of each ligand (Figs. 2E and 2F). We further performed CD spectroscopy analysis of four aporphine ligands to determine that 14148 is racemic and 15856 is an *R* stereoisomer (Supplementary Fig. 7).

To confirm specific interaction of each compound with 5-HT_2C_, we incubated individual compounds with the purified receptor and performed affinity MS binding assay. All four compounds showed significant binding to 5-HT_2C_ relative to the control (Fig. 2G). Furthermore, in a ligand competition experiment, all four compounds substantially reduced the binding of reference agonist and antagonist to 5-HT_2C_ in a concentration-dependent manner (Figs. 2H and 2I).

### Discovery of a 5-HT_2C_ selective and G_q_ biased agonist

We subsequently assessed the four aporphines for their binding affinity to three 5-HT_2_ subtypes using a radiolabeled ligand displacement assay. All four ligands displayed binding affinities to the three receptors with K_i_ values in the medium to high nanomolar range, and showed no significant preference of binding to 5-HT_2C_ (Figs. 3A-C, Supplementary Table 4). Intriguingly, 1857 and 15781 possess exquisite subtype selectivity for 5-HT_2C_ in G_q_-mediated calcium signaling: both are partial agonists (1857 EC_50_=308 nM, E_max_=86.1%; 15781 EC_50_=653 nM, E_max_=65.6% relative to 5-HT) whereas none of them exhibited measurable G_q_ agonism for 5-HT_2A_ or 5-HT_2B_ (Figs. 3D and 3E). In fact, both are weak antagonists of 5-HT_2A_ or 5-HT_2B_ (IC50 > 10 µM) (Supplementary Figs. 8A and 8B). In contrast to their remarkable selectivity for 5-HT_2C_, the approved anti-obesity drug lorcaserin only displayed moderate or no selectivity for 5-HT_2C_ over 5-HT_2B_ or 5-HT_2A_ (Fig. 3F). The other two aporphines, 14148 and 15856, are both non-selective antagonists of G_q_-mediated activity with submicromolar potency at three 5-HT_2_ subtypes (Supplementary Figs. 8C and 8D). It is worth noting that 1857 (agonist) and 14148 (antagonist) only differ by a single methyl substituent, yet they displayed opposite activity at 5-HT_2C_.

**Figure 3.**
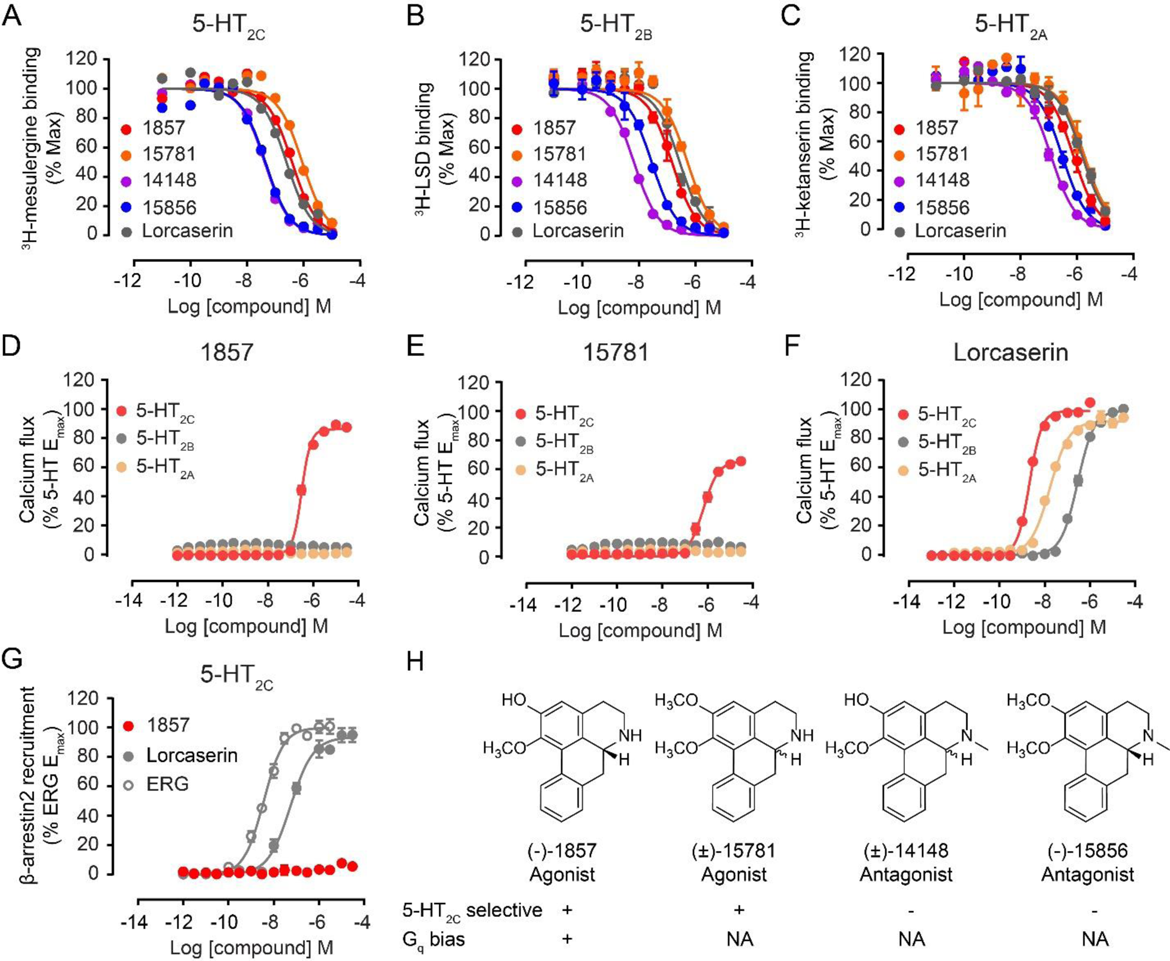
Pharmacological profiles of aporphine ligands. (A-C) Radiolabeled ligand binding curves for 5-HT_2C_, 5-HT_2B_ and 5-HT_2A_ in the presence of aporphines or lorcaserin. See also Supplementary Table 4 for K_i_ values. (D-F) G_q_-mediated calcium mobilization induced by 1857, 15781 and lorcaserin. 1857 and 15781 displayed partial agonism only at 5-HT_2C_ while lorcaserin activated three 5-HT_2_ receptors. (G) β-arrestin2 recruitment stimulated by 1857, locaserin and ergotamine (ERG, a known β-arrestin2 biased agonist for 5-HT_2C_). 1857 has no measureable agonist activity following ERG treatment. (H) SFSR summary of aporphines discovered in this study. *N*-unsubstituted aporphine 1857 and 15781 act as selective 5-HT_2C_ agonists. 1857 also displays G_q_ bias with no measurable β-arrestin activity. *N*-methyl substituted aporphine 14148 and 15856 act as non-selective 5-HT_2_ antagonists. NA, not assayed. Data represent means ± SEM of three independent experiments performed in triplicate.

For the partial agonist 1857, we then assessed its ability to recruit β-arrestin2 to 5-HT_2C_ using a reporter gene-based Tango assay^50^. Strikingly, 1857 exhibited no β-arrestin recruitment activity (Fig. 3G). In fact, it acts as an antagonist of 5-HT_2C_ related β-arrestin signaling against agonist ergotamine (Supplementary Fig. 8E). We also employed a bioluminescent resonance energy transfer (BRET) orthologous method that measures β-arrestin2 association with the receptor. This assay revealed that 1857 does not possess agonist activity in β-arrestin2 recruitment but acts as an antagonist for β-arrestin2 association (Supplementary Fig. 8F), confirming its G protein bias. In contrast, lorcaserin functioned as a full agonist at 5-HT_2C_ in both G protein and β-arrestin pathways (Figs. 3F and 3G).

In summary, 1857 displays a very unique pharmacological profile as a highly selective partial agonist for 5-HT_2C_ with an exclusive bias towards G_q_ signaling. Moreover, our results uncovered 5-HT_2C_ structure–functional selectivity relationship (SFSR) for a group of aporphine alkaloids which demonstrate either subtype-selective G protein preference or nonselective antagonism, dependent on *N*-6 substitution (Fig. 3H).

### Structural model of 5-HT_2C_ activation by 1857

We next predicted the binding pose of 1857 based on the solved 5-HT_2C_ crystal structure^15^ (PDB ID: 6BQG) by molecular docking. Embedded deep in the orthosteric pocket of the receptor, 1857 forms a salt bridge between its protonated nitrogen and the conserved aspartate D134^3.32^ which is a key interaction conserved in 5-HT and other aminergic GPCRs^13–15^ (Fig. 4A). In addition, 1857 forms extensive interactions with residues on transmembrane (TM) helices 3, 5, 6 and 7. Especially, the aporphine rings form π-π interactions with both F327^6.51^ and F328^6.52^ and hydrophobic interactions with V135^3.33^, A222^5.46^ and V354^7.39^ (Fig. 4A). In contrast, lorcaserin, which is smaller in size than 1857, leaves more space in the pocket and makes less extensive interactions with the aforementioned residues (Fig. 4B).

**Figure 4.**
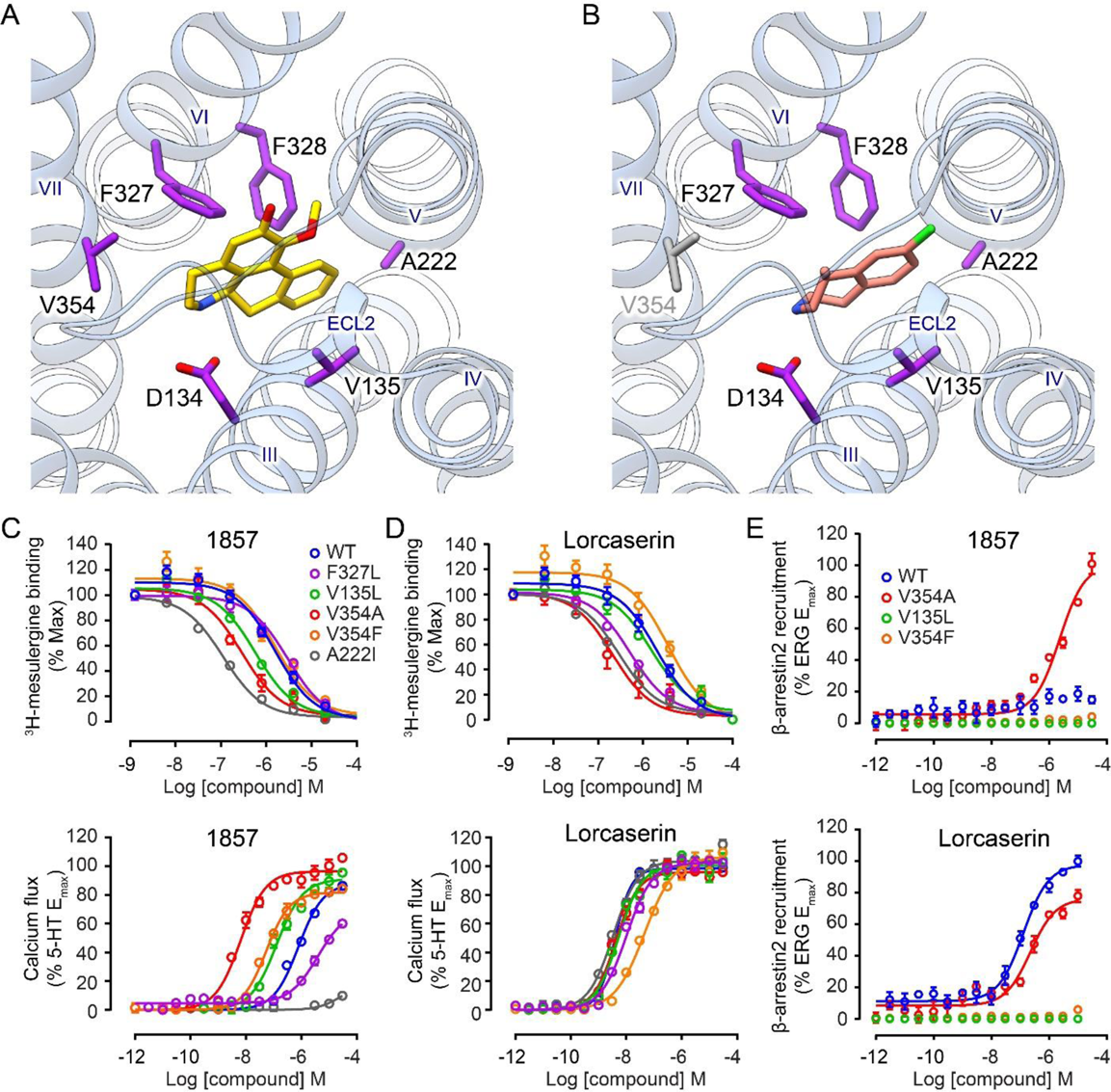
Key interactions between 5-HT_2C_ and 1857 specifically modulate the agonist activity. Docking poses of 1857 (A) and lorcaserin (B) in 5-HT_2C_. Predicted key interacting residues are in purple. V354^7.39^ in grey (B) does not interact with lorcaserin. Radiolabeled ligand binding curves (upper) and G_q_-mediated calcium flux (lower) in cells expressing wild-type (WT) or mutant 5-HT_2C_ in the presence of 1857 (C) or lorcaserin (D). (E) β-arrestin2 recruitment in cells expressing WT or mutant 5-HT_2C_ in the presence of 1857 (upper) or locaserin (lower). G_q_ activity of 5-HT_2C_ elicited by 1857 was significantly affected by mutations on five key interaction sites relative to WT yet they hardly changed G_q_ activity of 5-HT_2C_ treated by lorcaserin. See also Supplementary Table 5 for IC50/EC_50_ values. Data represent means ± SEM of three independent experiments performed in triplicate.

Mutating F327 to L to impair the predicted π-π interaction attenuated 1857’s binding affinity and G_q_-mediated agonist activity (Fig. 4C), yet this mutation did not affect lorcaserin’s affinity or activity (Fig. 4D). Another mutation V135L which may strengthen the hydrophobic interaction with the ligand significantly increased 1857’s affinity and promoted its G_q_-mediated agonist activity, whereas slightly increased affinity and no change of G_q_ activity was observed on lorcaserin (Figs. 4C and 4D). Most strikingly, V354A showed ∼300-fold increase of G_q_ activity (EC_50_=6.4 nM) though its binding was only enhanced by 6-fold (Fig. 4C, Supplementary Table 5). Mutating V354 to a bulky residue F also resulted in substantially increased G_q_ activity (EC_50_=54.0 nM) but affinity remained unchanged (Fig. 4C, Supplementary Table 5). It seems that V354 in TM7 is a critical residue specifically influencing G_q_-coupled activation by 1857 but not lorcaserin (Fig. 4D). Furthermore, among the mutants with increased G_q_ activity, 1857 only induced weak β-arrestin recruitment activity of V354A (EC_50_=2.3 μM) while the others stayed inactive in this pathway (Fig. 4E). Finally, A222I substantially increased 1857’s affinity possibly due to its longer aliphatic side chain. However, this mutation almost abolished G_q_ activity elicited by 1857, which may be related to a locked conformation of TM5 as a result of the strengthened hydrophobic interaction (Fig. 4C). Taken together, our docking supported by mutagenesis study pinpointed key interactions specific for G_q_-coupled 5-HT_2C_ activation by 1857 and not by lorcaserin.

### Agonist 1857 suppresses food intake and induces weight loss in mice

Prior to evaluating 1857 in animal models, we measured its brain permeability by intravenous injection to mice (10 mg/kg; Supplementary Table 6). Brain-to-plasma ratio of 1857 stayed as high as 6.4 at 4 h after dosing, indicating its excellent pharmacokinetics profile. Since 5-HT_2C_ is a potential therapeutic target for obesity, we investigated the effects of 1857 on food intake in comparison with lorcaserin. In the first experiment, acute intraperitoneal administration of 1857 (30 mg/kg) reduced food intake from 30 min to 4 h post-injection (*P* <0.01) whereas lorcaserin (10 mg/kg) suppressed food intake up to 2 h post-injection (*P* <0.05) (Figs. 5A and 5B). Thus, 1857 administered at three times the dose of lorcaserin produced very similar *in vivo* efficacy, despite that its *in vitro* potency for 5-HT_2C_ activation is two orders of magnitude lower than that of lorcaserin (Figs. 3D and 3F). This result could be attributed to better brain permeability and metabolic stability of 1857.

We then examined the effect of 1857 in diet-induced obesity (DIO) mice. Mice were fed with high fat diet for 2 months to induce obesity before receiving daily treatment of 1857 (30 mg/kg). Significant reduction of food intake (Fig. 5C) and weight loss (Fig. 5D) started to be observed in mice following treatment for 5 days and lasted till the end of this study. Consistent with weight loss, 1857 treated DIO mice had lower blood glucose (−20.9%), reduced serum total cholesterol (TC, −7.9%) and unchanged serum total triglyceride (TG) relative to vehicle (Fig. 5E). Furthermore, 1857 treatment also decreased liver TC and TG levels (Fig. 5F) as well as the total organ weight of epididymal white adipose tissue (WAT, −29.3%) and inguinal WAT (−46.4%) in DIO mice (Fig. 5G). Similar effects were reported in DIO rats treated with lorcaserin^51^. Therefore, our study confirmed the efficacy of 1857 in attenuating obesity which is in line with its specific activity on 5-HT_2C_.

**Figure 5.**
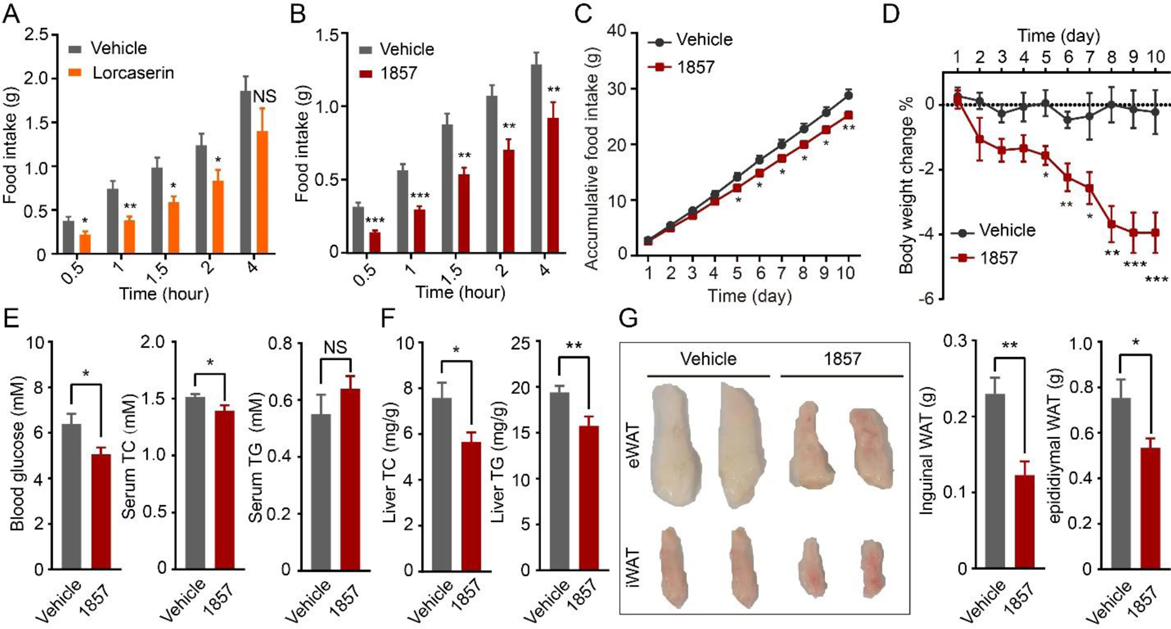
*In vivo* anti-obesity effects of 1857. Acute food intake suppression induced by lorcaserin (A) or 1857 (B). Overnight fasting mice (n=8 each group) were treated with lorcaserin (10 mg/kg), 1857 (30 mg/kg) or vehicle 30 min before feeding. Food intake was measured at indicated time points. (C-G) 1857 inhibited food intake and showed anti-obesity effects in a diet-induced obesity (DIO) mouse model. DIO mice (n=9 each group) were treated with 1857 (30 mg/kg) or vehicle daily for 10 days. Accumulative food intake (C) and body weight change (D) were recorded during the treatment. Blood and liver were collected to measure blood glucose, serum cholesterol (TC) and triglyceride (TG) levels (E) as well as liver TC and TG levels (F). The weight of white adipose tissues (WAT) was also measured and representative tissue images were shown (G). Data represent means ± SEM. \**P* <0.05, \*\**P* <0.01 and \*\*\**P* <0.001 (two-tailed Student’s *t*-test).

## Discussion

In this study, we established an affinity MS-based approach specifically for GPCR ligand discovery from natural products. To identify both agonists and antagonists for a given GPCR, the ensemble of the purified receptor used as a bait for ligand enrichment is ought to comprise both active and inactive conformations. However, a large number of GPCR constructs optimized for *in vitro* purification and structural characterization tend to yield proteins predominantly at inactive conformational states due to the inherent flexibility and instability of functionally active states^52–56^. Therefore, previous affinity MS screens of synthetic compound libraries with a purified GPCR only discovered new antagonists^45, 57^ that readily bind to a receptor in the inactive state. To drive the heterogeneous population of 5-HT_2C_ towards active conformations, we reversed the thermostabilizing mutation C360N^7.45^ originally designed for receptor crystallography^15^, which increased our chances of capturing agonists. In another study of using GPCR-expressing cell membranes for affinity MS screen, we also observed that all agonizing ligands identified with a wild-type GPCR abrogated their binding to the receptor when it incorporated multiple thermostabilizing mutations^44^. Therefore, careful design of the construct to shift the receptor conformation to the active state is essential for agonist discovery using this approach.

The naturally occurring compounds 1857 ((*R*)-asimilobine) and 15781 (nornuciferine) discovered here possess an aporphine scaffold which represents a novel chemotype for 5-HT_2C_ agonists. Compounds in this alkaloid subclass have been traditionally characterized as ligands for the dopamine receptor system and play potential therapeutic roles in the treatment of Parkinson’s disease and other neurological disorders^58–60^. With regard to serotonin receptors, aporphines have mostly been studied as ligands for 5-HT1A and 5-HT_2A_ receptors^48^ and are rarely associated with 5-HT_2C_. Notably, the majority of natural or synthetic aporphines exhibit antagonistic activities at serotonin receptors with no documented subtype selectivity or signaling bias^48, 49, 61^. In contrast, 1857 is the first aporphine displaying exclusive biased G protein agonism at 5-HT_2C_ with exquisite selectivity over 5-HT_2A_ and 5-HT_2B_. The unique pharmacological profile of 1857 opens a new avenue for design of potent and functionally selective 5-HT_2C_ ligands with great potential in the treatment of obesity, schizophrenia and other neurological disorders^26^. Moreover, 1857 could serve as a desirable probe for elucidating the structural basis of preferential G protein signaling (details in Supplemental Note 3). Among the four aporphine ligands discovered in this study, 1857 and 15858 isolated from herbs are optically pure stereoisomers while the other two aporphines (15781, 14148) are racemic (Fig. 3H). How the stereospecificity of different ligands affects the pharmacological properties awaits further investigation. In addition, although 1857 shows remarkable G_q_-coupling selectivity at 5-HT_2C_, its broader selectivity among the 5-HT receptor subfamily and other aminergic GPCRs remains to be determined.

The natural product screening approach presented in this study enables rapid discovery of GPCR ligands with sophisticated pharmacological properties. Although we identified a new series of aporphine ligands for 5-HT_2C_, the rest 95 hits from screening ST extract fractions may contain more 5-HT_2C_ modulators with novel structures and distinct bioactivities (Fig. 2C, Supplementary Table 2). This high-throughput and unbiased screening approach which is generalizable to other receptors would accelerate the exploration of largely untapped natural product chemical space for discovering novel and improved drug leads targeting GPCRs.

## Methods

### Receptor expression and purification

The protein expression and purification has been described previously^15, 61^. In brief, the 5-HT_2C_R-BRIL construct was subcloned into a modified pFastBac1 vector (Invitrogen). The construct was optimized with the truncation of N-terminal residues (1-39) and C-terminal residues (393-458). The residues of third intracellular loop (IL3) from L246 to M300 were replaced by thermostabilized apocytochrome b562RIL (BRIL). No mutation sites were incorporated into this construct. The construct was then expressed in *Spodoptera frugiperda* (*Sf9*) cells with a haemagglutinin (HA) tag followed by a FLAG tag at the N terminus and a 10× His tag at the C terminus. Cells were cultured at 27 °C and harvested after 48 h post infection. 5-HT_2C_R-BRIL membrane preparation and protein purification were performed with the same procedure as mentioned before^15, 61^. The purity and monodispersity of the 5-HT_2C_ protein were measured by analytical size-exclusion chromatography (aSEC).

### Herbal extract preparation

Nine herbs examined in this study are Aristolochia debilis (AD), Tetradium ruticarpum (TR), Rauvolfia verticillata (RV), Strychnos nux-vomica (SN), Stephania tetrandra (ST), Ziziphus jujubam (ZJ), Menispermum dauricum (MD), and Catharanthus roseus (CR). Each of them was first pulverized into powder. The powder (200 g) was extracted with 500 mL of 70% ethanol by waterbath ultrasonication for 30 min. The extraction was performed three times in total. Then the organic solvent was removed by vacuum evaporation at 50 °C. The residual material was dissolved in 0.3% (v/v) hydrochloric acid and partitioned with EtOAc three times. Then the aqueous layer was basified with 5% (v/v) ammonia to pH 9-10 and partitioned with EtOAc three times. The EtOAc phase was dried out and the powder was stored at −80 °C. The stock solution (100 mg/mL) of each herbal crude extract was prepared by dissolving the powder with 95% DMSO and was stored at −20 °C.

### Affinity MS screening of herbal extracts

The method developed for GPCR ligand screening from compound libraries^45^ was adapted to herbal extract screening here. The purified protein 5-HT_2C_ or HCA2 (3 μg) was immobilized on nickel agarose beads (Sigma) in the incubation buffer containing 50 mM HEPES, pH 7.5, 150 mM NaCl, 0.05% (w/v) DDM, 0.01% (w/v) CHS at 4 °C overnight. The stock of each herbal extract was diluted with the incubation buffer to a final concentration of 0.5 mg/mL. Then the 5-HT_2C_ beads were incubated with the diluted crude extract at 4 °C for 1 h. The supernatant was removed and the beads were washed four times with 150 mM ammonium acetate (pH 7.5) after incubation. The compounds bound to 5-HT_2C_ were then dissociated with 200 μL methanol, dried out in a speed vacuum, and reconstituted in 50% methanol before LC-MS/MS analysis. The control sample was prepared with the same procedure by using HCA2 beads. All samples were prepared in four independent replicates. Samples were analyzed on a Shimazu L30A UPLC system (Shimazu) coupled to a TripleTOF 6600 mass spectrometer (AB SCIEX) operating in the positive ion mode. Chromatographic separation was performed on a ZORBAX Eclipse Plus C18 column (3.5 µm, 2.1×100mm, Agilent) at a flow rate of 300 μL/min and maintained at 40 °C with the mobile phases of water/0.1% formic acid (A) and acetonitrile/0.1% formic acid (B). The LC gradient was as follows: 0−2 min, B at 5%; 2−12 min, B at 10%−30%; 12−25 min, B at 30%-90%; 25−30 min, B at 90%-90%, then re-equilibrate for 5 min. Full-scan mass spectra were acquired in the range of 100-1500 m/z with major ESI source settings: voltage 5.0-5.5 kv; gas temperature 500°C; curtain gas 35 psi; nebulizer gas 55 psi; and heater gas 55 psi. MSMS spectra were acquired on top 10 precursors with collision energy set at 45 eV with a CE spread of 15 eV and other ion source conditions identical to the full scan.

### Metabolomics data processing for 5-HT_2C_ ligand identification

Compounds in the target and control samples were identified by extracting ion chromatograms (EICs) using Peakview 2.2 (AB SCIEX) based on accurate mass (<10 ppm), isotope envelop matching (<10% deviation) in accordance with the compound formula registered in TCMHD and retention time (<0.2 min) matched with peaks detected in the crude extract. Binding index (BI) of each compound is defined to be the ratio of MS intensity of the compound detected in the 5-HT_2C_ target vs control. Initial hits were selected based on a mean BI >2 and *P* <0.05 from four experimental replicates. Significant difference of each compound’s MS intensity between target and control samples was determined by a two-tailed *t*-test with Bonferroni correction.

### Stephania tetrandra (ST) crude extract fractionation

The powder of the ST crude extract (200 mg) was dissolved in methanol and then fractionated using a Sunfire C18 OBD Prep column (5 µm, 19×250mm, Waters) running at a flow rate of 10 mL/min with the mobile phase of water/0.1% formic acid (A) and acetonitrile (B). The LC gradient was 0−3 min, B at 10%; 3−20 min, B at 10%−90%; 20−25 min, B at 90%-90%. Three fractions were collected according to the UV response and LC separation. The solvent was removed by vacuum evaporation at 50 °C and the residue was stored at −80 °C. Each fraction was reconstituted in the same incubation buffer to the same concentration as described for the ST crude extract before the affinity MS screen.

### Isolation of two putative ligands from the ST extract

Air-dried, powdered roots of *Stephania tetrandra* (3 kg) were extracted three times with 95% ethanol at room temperature. Then the organic solvent was removed and the residue underwent the same procedure as described in the natural herb extract preparation. The crude extract (100 g) was separated on a silica gel column and eluted with CHCl3-MeOH (1:0-1:1) to obtain three fractions (F1-F3). Each fraction was then analyzed with UPLC-DAD/MS to identify the putative ligands from affinity MS screen. In Fraction F3, detection of two peaks at *m/z* 268. 1334 and 282.1488 which showed the same retention time as ligands 1857 and 15781 indicated the presence of the two expected aporphine alkaloids. Compounds 1857 (2.1 mg) and 15781 (10.2 mg) were then purified from Fraction F3 by Shimadzu LC-20A (Shimadzu) using Sunfire C18 column (5 µm, 19×250mm, Waters) with the gradient ACN-H2O (20-50%) at a flow rate of 10 mL/min. The peaks at 8.9 min and 12.1 min (for 1857 and 15781 respectively) were collected separately and dried by a vacuum evaporation. The structures of the two pure compounds were elucidated with 1D (^1^H and ^13^C) and 2D (HSQC and HMBC) NMR (Avance III HD 800 MHz, Bruker) analysis and data are shown in the Supporting Information. The configurations of the two compounds were further determined based on their optical rotation values measured with an automatic polarimeter (Autopol VI, Rudolph).

### Affinity MS-based validation of pure ligand binding to 5-HT_2C_

Compounds 14148 and 15856 were purchased from BioBioPha Co. Ltd (Kunming, China) and Chengdu Herbpurify Co. Ltd (Chengdu, China), respectively. Compounds 1857 and 15781 were in-house isolated as described above. Their structures are described in Table S3. These four aporphine alkaloids were mixed at a final concentration of 100 nM for each compound. Then the compound mixture was incubated with purified 5-HT_2C_ or HCA2 proteins under the same conditions of the ligand screening experiment. The receptor-associated compounds were analyzed by LC-MS/MS. In the affinity MS binding assay of this simple mixture, a short LC gradient was applied for compound separation: 0−2 min, B at 5%; 2−2.1 min, B at 5%−10%; 2.1−5 min, B at 10%-30%; 5−5.1 min, B at 30%-90%; 5.1−7 min, B at 90%.

Specific compound peaks in target and control samples were extracted using PeakView 2.2 (AB SCIEX) based on the accurate mass measurement (<10 ppm) and RT matching with the standard (<0.2 min). BI and *P* values (n = 4) were determined for each compound as described in the herbal extract screening experiment to assess ligand binding specificity for 5-HT_2C_.

### Affinity MS-based ligand competition assay for 5-HT_2C_

The 5-HT_2C_ agonist 5-MeO-DMT and antagonist ritanserin were used as marker ligands in the ligand competition assay. Purified 5-HT_2C_ protein immobilized on nickel agarose beads (Sigma) was incubated with a given marker ligand at 2 nM mixed with each test compound (aporphine ligands) at an increased concentration (0, 2, 20 μM) at 4 °C for 1 h. After incubation, the compounds were dissociated from the 5-HT_2C_ and analyzed by LC-MS/MS using the same method in the previous pure ligand binding assay. The MS intensity of the marker ligand under different conditions was extracted from the raw data using PeakView 2.2 (AB SCIEX) based on the same criteria described in the previous pure ligand binding assay. Reduction of the marker ligand response indicated the extent of binding competition by each test compound. Two independent experiments were performed in technical duplicate under each condition.

### Cloning and mutagenesis

Mutagenesis of the 5-HT_2C_ construct was performed according to the Q5® site-Directed Mutagenesis Kit protocol (New England BioLabs). In brief, PCR reactions were performed using the wild-type 5-HT_2C_ receptor cDNA (pcDNA3.1) and primers containing the mutation sites of interest to create mutant plasmids. After DpnI (New England BioLabs) digestion of the parental DNA and transformation, positive clones were selected by ampicillin resistance. DNA was prepared using Miniprep Kit (Axygen) and sequenced (Genewiz) using forward (CMV) and reverse (BGHreverse) sequence primers.

### Radioligand binding assay

Radioligand binding assays for wild-type receptors were performed using membranes prepared from 5-HT_2A_/2B/2C transfected HEK293 cell lines. Radioligands used in the assays were ^3^H-ketansetin (PerkinElmer; specific activity = 42.5-47.3 Ci/mmol) for 5-HT_2A_; ^3^H-LSD (PerkinElmer; specific activity = 82.9-83.3 Ci/mmol) for 5-HT_2B_; ^3^H-mesulergine (PerkinElmer; specific activity = 80.9-83.0 Ci/mmol) for 5-HT_2C_. The unlabeled ligands were prepared in binding buffer (50 mM Tris, 10 mM MgCl2, 0.1 mM EDTA, 0.1% BSA, 0.01% ascorbic acid, pH 7.4) ranging from 10 μM to 30 pM. Assay plates were incubated at room temperature for 1 h and then they were harvested using vacuum filtration onto 0.3% polyethyleneimine-presoaked 96-well filter mats A (PerkinElmer) and washed three times with cold wash buffer (50 mM Tris, pH 7.4). Scintillation cocktail (Meltilex) was melted onto the dried filters and then the plate was read using a Wallac Trilux Microbeta counter (Perkin Elmer). Data were analyzed with GraphPad Prism 7.0 (Graphpad Software Inc.) using ‘one site-Fit K_i_’ to obtain K_i_. Data were normalized to the top (100%, no competitor) and bottom (0%, 10 µM clozapine for 5-HT_2A_, 10 µM SB206553 for 5-HT_2B_, 10 µM ritanserin for 5-HT_2C_) to represent the percent of displacement. These assays were conducted by NIMH PDSP, directed by Bryan L Roth, M.D., Ph.D., the University of North Carolina at Chapel Hill, North Carolina, and Program Officer Jamie Driscoll at NIMH, Bethesda, MD.

To assay the mutants, they were first established by PCR-based site-directed mutagenesis and confirmed by DNA sequencing. The wild-type or mutant 5-HT_2C_ was cloned into the pcDNA3.1 vector (Invitrogen) for transfection. CHO cells were washed twice 24 h after transfections and incubated with blocking buffer (F12 supplemented with 33 mM HEPES and 0.1% bovine serum albumin (BSA), pH 7.4) for 2 h at 37 °C. Subsequently, the cells were incubated in binding buffer (DMEM supplemented with 25mM HEPES and 0.1% BSA) with a constant concentration of ^3^H-mesulergine (1 nM) and different concentrations of unlabeled 5-HT (1.28 nM ∼ 100 μM), lorcaserin and 1857 (0.64 nM ∼ 50 μM) at room temperature for 3 h. Cells were washed three times with ice-cold PBS and lysed by 50 μl lysis buffer (PBS supplemented with 20 mM Tris-HCl, 1% Triton X-100, pH 7.4). The plates were subsequently counted for radioactivity (counts per minute, CPM) in a scintillation counter (MicroBeta2 Plate Counter, PerkinElmer) using a scintillation cocktail (OptiPhase SuperMix, PerkinElmer).

### Calcium mobilization assay

HEK293T cells stably transfected with wild-type 5-HT_2A_, 5-HT_2B_ or 5-HT_2C_ were plated into poly-lysine coated 384-well black clear bottom plates at a density of 15,000 cells per well with 40 μL DMEM with 1% dialyzed FBS. For mutant activity evaluation, T-rex 293 cells (approximately 3 × 10^6^ cells per 10-cm dish) were transfected with 5-HT_2C_ wild-type or mutant DNA using Lipofectamine 2000 (Invitrogen) following manufacturer’s protocols. After 18-24 h transfection, cells were plated at the same condition described above for stable cells. Next day, the media was decanted and cells were incubated for 1 h at 37 °C with Fluo-4 Direct dye (Invitrogen) in FLIPR buffer (1× HBSS, 2.5 mM probenecid, and 20 mM HEPES, pH 7.4). After the dye loaded, the cells were placed in a FLIPR^TETRA^ fluorescence imaging plate reader (Molecular Devices). Drugs were diluted at 3× final concentration in drug buffer (1× HBSS, 0.1% BSA, 20 mM HEPES, pH 7.4) and aliquotted into 384-well plates, which were then placed in the FLIPR^TETRA^. The fluidics module and plate reader of the FLIPR^TETRA^ were programmed to read baseline fluorescence for 10 s (1 read/s), then to add 10 μL of drug dilutions per well and to read for 3 min (1 read/s). Fluorescence of each well was normalized to the average of first 10 reads. Then the maximum-fold increase, which occurred within 60 s after drug addition, over baseline fluorescence elicited by vehicle or drug was determined. For the antagonist mode, 5-HT (3 nM) was used to activate the receptor. Data were normalized to percent 5-HT simulation and EC_50_ or IC50 was analyzed in GraphPad Prism 7.0 (Graphpad) using log (agonist) vs response or log (antagonist) vs response.

### Tango β-arrestin2 recruitment assay

The Tango constructs of human 5-HT_2C_ and its mutants were designed and β-arrestin2 recruitment assay was performed as described previously^50^. Briefly, HTLA cells were transfected with 5-HT_2C_ Tango plasmid or mutants. After at least 24 h, HTLA cells were plated into poly-lysine coated 384-well white clear bottom plates at a density of 15,000 cells per well with 40 μL DMEM containing 1% dialyzed FBS. Six hours later, cells were simulated with 20 μL per well drug dilutions (3 ×) prepared in drug buffer (1× HBSS, 0.1% BSA, 20 mM HEPES, pH 7.4) and incubated at 37 °C overnight. Then media and drug solutions were decanted and 20 μL per well of BrightGlo reagents (Promega) were added. After 20 min incubation, luminescence was read on an Envision counter (Perkin Elmer). For the antagonist mode, agonist ERG EC80 (50 nM) was used to activate the receptor. Data were normalized to percent ERG simulation and EC_50_ or IC50 was analyzed in GraphPad Prism 7.0 (Graphpad) using log (agonist) vs response or log (antagonist) vs response.

### BRET β-arrestin2 recruitment assay

We measured the effect of 1857 on the recruitment of β-arrestin2 in CHO cells stably expressing 5-HT_2C_-Rluc8 and β-arrestin2-Venus by bioluminescence resonance energy transfer (BRET) assay. The cells were seeded onto 96-well plate at a density of 3 × 10^4^ cells per well. Prior to BRET experiments, cells were rinsed twice with HBSS and then incubated with fresh HBSS for 30 min at 37 °C. After incubation with various concentrations of 1857 for 15 min, 5 μM coelenterazine-H (ThermoFisher) were added followed by 5 min incubation. Base-line BRET signals were read immediately at 470 nm and 535 nm for 11 cycles using an EnVision instrument (PerkinElmer). Constant concentration of the agonist lorcaserin (4 µM) was then added and detected for another 49 cycles. Data are presented as a BRET ratio, calculated as the ratio of Venus to Rluc8 signals after subtracting the lorcaserin value.

### Prediction of ligand binding poses by molecular docking

Molecular docking was performed with Scrodinger Suite 2015-4. Crystal structure of 5-HT_2C_ with agonist ergotamine (PDB ID: 6BQG)^15^ was used. Processing of the protein structure was performed with the ‘Protein Preparation Wizard’. Converting of ligands from 2D to 3D structures was performed using ‘LigPrep’. Molecular docking was performed with Glide 6.9 in standard precision.

### Acute feeding suppression

Male C57BL/6J mice were allowed to habituate to single cage housing and daily intraperitoneal injection of saline was given one week before starting the experiments (lights on/off 0700/1900). Food was removed at 19:00 for overnight fasting, the mice (N = 8 each group) were then treated with vehicle, lorcaserin (10 mg/kg), or 1857 (30 mg/kg) at 9:00 the next day. Food was returned 30 min there after. Food intake was measured at 30, 60, 90, 120, 240 min after the presentation of food. Data represented are means ± SEM. Significant differences among groups were determined by unpaired, two-tailed Student’s *t*-test.

### DIO mice feeding suppression

Male C57BL/6J mice were fed with high fat diet (HFD, 60% of energy from fat, Research Diets, no. D12492) for 8-9 weeks. The mice were housed individually and allowed *ad libitum* access to water and fed with HFD. For the subchronic study (10-day treatment), DIO mice (N = 9 each group) were i.p. injected with vehicle or 1857 (30 mg/kg) daily. The body weight and food intake were monitored daily. At the end of the study, the mice were anesthetized, and blood, liver, and white adipose tissues (WAT) were collected for further analysis. Blood glucose was measured from the tail vein with a glucometer. Serum was prepared by centrifuging at 2000 × g for 10 min. Livers were homogenized in chloroform-methanol (2:1). The organic phase was further dried under N2 and then resolved in ethanol. Total cholesterol (TC) and total triglyceride (TG) in serum or extracted liver samples were measured with TC and TG kits (E1005-250, E1003-250, Applygen). TC and TG levels in the liver samples were normalized to the liver weight. Data represented are means ± SEM. Significant differences among groups were determined by unpaired, two-tailed Student’s *t*-test.

## Acknowledgements

We thank Dr. Raymond C. Stevens for valuable advice, Dr. Shuguang Yuan and Dr. Chu Wang (Peking University) for fruitful discussion. We also thank Xiaoyan Liu, Junlin Liu and Fangfang Zhou from iHuman Institute, Antao Dai and Chao Zhang from Shanghai Institute of Materia Medica for technical assistance. This work was funded by ShanghaiTech University, National Key R&D Program of China grants (2018YFA0507000 (M.-W.W.), 2018YFA0507004 (W.S.), 2016YFC0905900 (G.Z.), 2017YFC1001300 (G.Z.)), National Mega R&D Program for Drug Discovery grants (2018ZX09711002-002-005 (D.H.Y.), 2018ZX09735-001 (M.-W.W)), National Natural Science Foundation of China grants (31971362 (W.S.), 81773792 (D.H.Y.), 31771130 (G.Z.)), Novo Nordisk-CAS Research Fund grants 2017 and 2019 (D.H.Y.). We are also grateful to the staff members of animal facility at the National Facility for Protein Science in Shanghai (NFPS), Zhangjiang Lab, China for their technical support.

## Author contributions

B.Z. performed receptor purification, affinity MS ligand screen and binding validation experiments with the help of H.C.; B.Z. and S.Z. evaluated compound *in vivo* efficacy. B.Z. performed receptor mutagenesis and cell signaling assays, and isolated compounds with assistance from Y.X., N.Y. and Y.X.; D.Y. X.Q.C. and W.S. performed radiolabeled ligand binding and BRET assays; X.H. performed radiolabeled ligand binding assays on wild-type 5-HT_2_ receptors and edited the manuscript. Y.W. conducted molecular docking analysis supervised by S.Z. Y.P. helped with 5-HT_2C_ purification supervised by Z.-J.L.. G.Z. and M.-W.W. were involved in the overall project management and edited the manuscript. W.S. and B.Z. wrote the manuscript with edits and inputs from all authors. W.S. conceived and supervised the project.

1857 brain and plasma pharmacokinetics were evaluated after a single intravenous dose of 10 mg/kg in male C57/6BJ mice (n=3 for each time point). After 30 or 240 minutes (min), mice were anesthetized to obtain whole blood and brain samples. Plasma was prepared from sodium heparin-treated whole blood and separated by centrifugation. Plasma and brain samples were frozen and stored at −80°C until analysis. Compound concentration (Conc.) was determined using single reaction monitoring (SRM)-based MS analysis. Compound with a brain/plasma ratio of greater than 1 is considered to be able to cross the blood-brain barrier freely. Data presented are means ± SEM.

The SRM method is as follows: A Waters Acquity I-Class UPLC coupled with a Xevo TQ-S mass spectrometer was used for determination of each compound concentration. LC separation was performed on a Waters Acquity BEH C18 (2.1×50 mm, 1.7 μm) column with the gradient being 0-1.2 min, B at 5%; 1.2-1.5 min, B at 70-95%; then re-equilibrating for 5 min. Mobile phase was water/0.1% formic acid (A solvent) and acetonitrile/0.1% formic acid (B solvent). MS instrument operated in an MRM mode with positive electrospray ionization to monitor analyte transitions of m/z 268.068 > 190.976 and m/z 282.068 > 265.07 for measurement of compound 1857 and 15781, respectively.

## Supplementary Note 1

### Identification of known 5-HT_2C_ agonists from herbal extracts

The established affinity MS workflow was first applied to screening 5-HT_2C_ ligands from crude extracts of AD and TR that displayed the highest potency among all TCM herbs studied. The purified receptor was incubated with either extract and underwent the same affinity MS procedure as described above. Representative total ion chromatograms for the crude extract, 5-HT_2C_ target and control samples are shown in Supplementary Fig. 3A. A targeted metabolomics data mining strategy previously developed by us^1^ was implemented to process the affinity MS screening data for individual extracts. All LC-MS features that matched the peak characteristics of compounds registered in the TCM herb database (TCMHD)^2^ were assigned to be herbal constituents. To identify these putative ligands, we determined the binding index (BI) for each of 704 and 485 assigned constituents in AD and TR extracts, respectively (Supplementary Table 1). Screening hits were selected if their mean BI values were above 2.0 (*P* < 0.05, *n* = 4)^3, 4^. In the end, three and ten initial hits were identified from AD and TR extracts, respectively (Supplementary Fig. 3B).

Interestingly, serotonin, the natural ligand for all 5-HT family members, turned out to be a top-ranking hit with high BI values in the screening (Supplementary Fig. 3B, Supplementary Table 1). A serotonin analogue 5-methoxy-*N*, *N*-dimethyltryptamine (5-MeO-DMT) known as 5-HT_2C_ agonist^5^ was also identified from TR screening (Supplementary Fig. 3B). MSMS spectral matching and retention time consistency between the putative ligand in 5-HT_2C_ and pure standard further confirmed the structural identity of two compounds (Supplementary Fig. 3C). Of note is that neither serotonin nor 5-MeO-DMT has been reported to be present in these two herbs, suggesting that the affinity MS screen in combination with metabolomics data mining is a powerful tool to identify unknown bioactive constituents in natural products.

To examine how much of the total bioactivity of AD and TR is attributed to serotonin and 5-MeO-DMT, we determined the percentage of each compound in the herbal extract (Supplementary Fig. 4). According to their measured concentration in the extract and potency of the pure agonist (Supplementary Fig. 3D), we estimated to what extent serotonin or 5-MeO-DMT accounted for the total agonism of an extract (Supplementary Fig. 3E). It turned out that serotonin alone or serotonin combined with 5-MeO-DMT contributed to 100% of the overall 5-HT_2C_ activity of AD and TR (Supplementary Fig. 3E). As serotonin is present in certain plants for growth regulation and defense^6^, we speculate that serotonin and its analogue are also biosynthesized in AD and TR. When analyzing additional extracts from other herbs with strong 5-HT_2C_ activities (RV and SR), serotonin was also detected at 0.02%-0.05%, close to that of TR. Thus, these two herbs were abandoned for further experimentation. Taken together, affinity MS screen conveniently identified known agonists accountable for the bioactivity detected in specific herbs, thereby prompting us to change the focus to other medicinal plants.

## Supplementary Note 2

(*R*)-Asimilobine (1857).^1^H NMR (CD3OD) δ 8.34 (d, *J* = 7.9 Hz, 1H), 7.31-7.26 (m, 2H), 7.23 (td, *J* = 7.3, 1.3 Hz, 1H), 6.64 (s, 1H), 3.71 (dd, *J* = 14.1, 4.5 Hz, 1H), 3.58 (s, 3H), 3.31-3.29 (m, 1H), 2.92-3.00 (m, 2H), 2.85 (dd, *J* = 13.8, 4.5 Hz, 1H), 2.69-2.66 (m, 2H); ^13^C NMR (CD3OD) δ 149.3, 143.7, 135.9, 132.2, 129.1, 127.5, 127.5, 127.1, 127.0, 126.6, 125.9, 115.2, 59.0, 53.4, 42.5, 36.5, 27.8. HRMS calcd for C17H17NO2 ([M + H]): 268.1332; found, 268.1336. [α]D −92.0 (c 0.1, MeOH)^7–9^.

Nornuciferine (15781).^1^H NMR (CD3OD) δ 8.34 (d, *J* = 7.9 Hz, 1H), 7.34 (dd, *J* = 14.9, 7.3 Hz, 2H), 7.29 (t, *J* = 7.3 Hz, 1H), 6.88 (s, 1H), 4.15 (dd, *J* = 14.1, 4.1 Hz, 1H), 3.90 (s, 3H), 3.66 (s, 3H), 3.65-3.61 (m, 1H), 3.30-3.20 (m, 2H), 3.04 (dd, *J* = 13.6, 4.4 Hz, 1H), 3.00 (dd, *J* = 17.0, 4.0 Hz, 1H), 2.89 (t, *J* = 13.6 Hz, 1H); ^13^C NMR (CD3OD) δ 153.4, 145.7, 133.7, 131.4, 128.2, 127.8, 127.7, 127.2, 126.8, 126.3, 123.5, 111.6, 59.2, 55.0, 53.0, 41.5, 34.3, 25.7. HRMS calcd for C18H20NO2 ([M + H]): 282.1489 m/z; found, 282.1491 m/z. [α]D − 1.5 (c 0.1, MeOH)^7^.

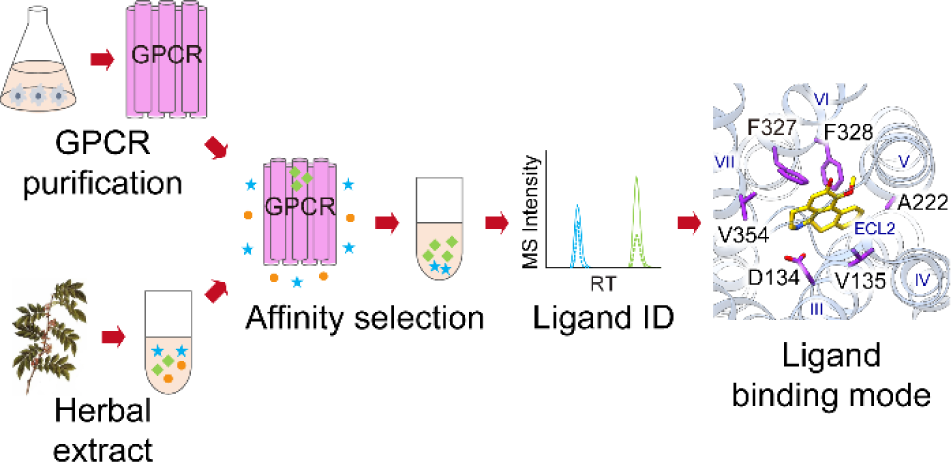

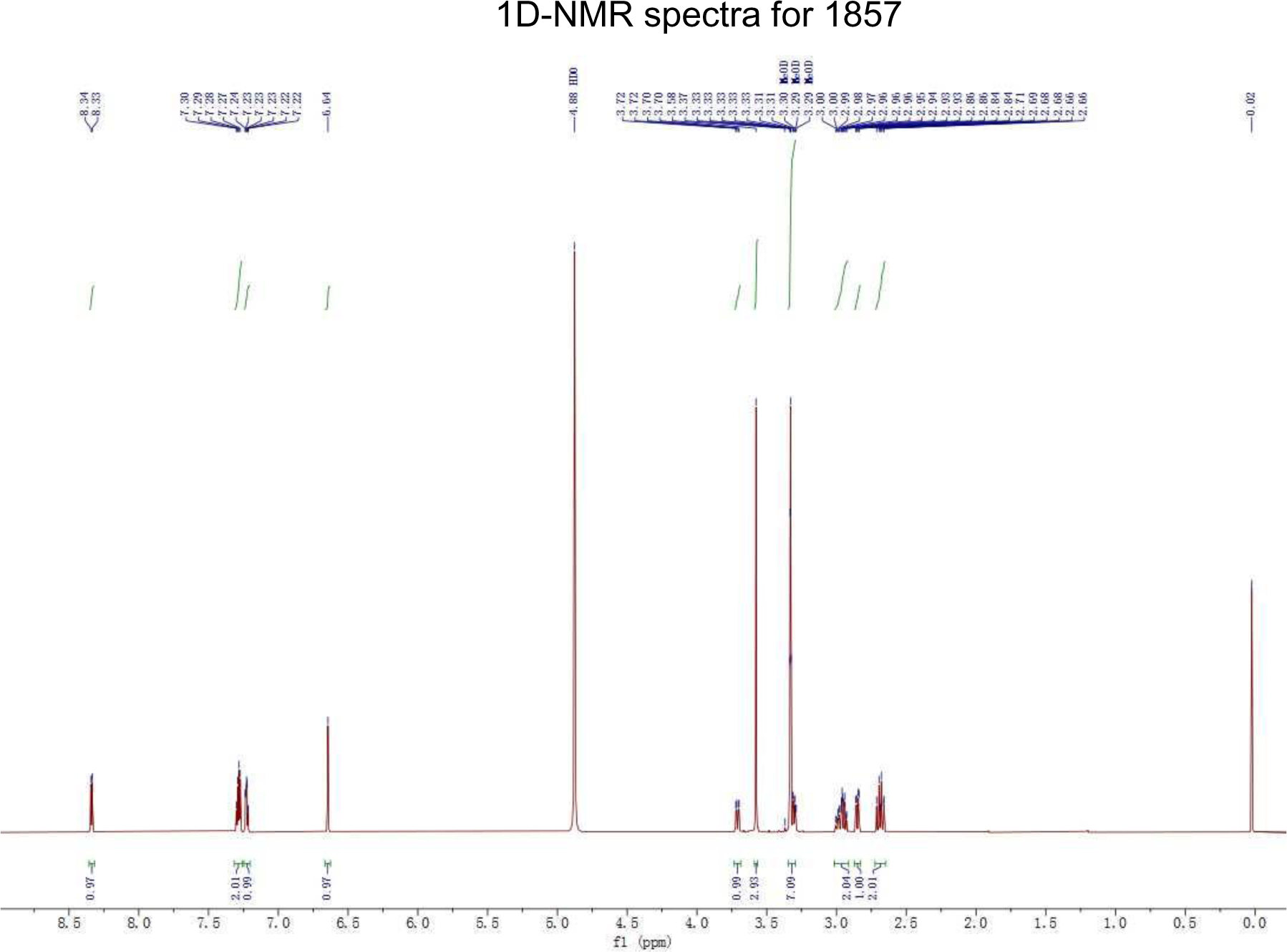
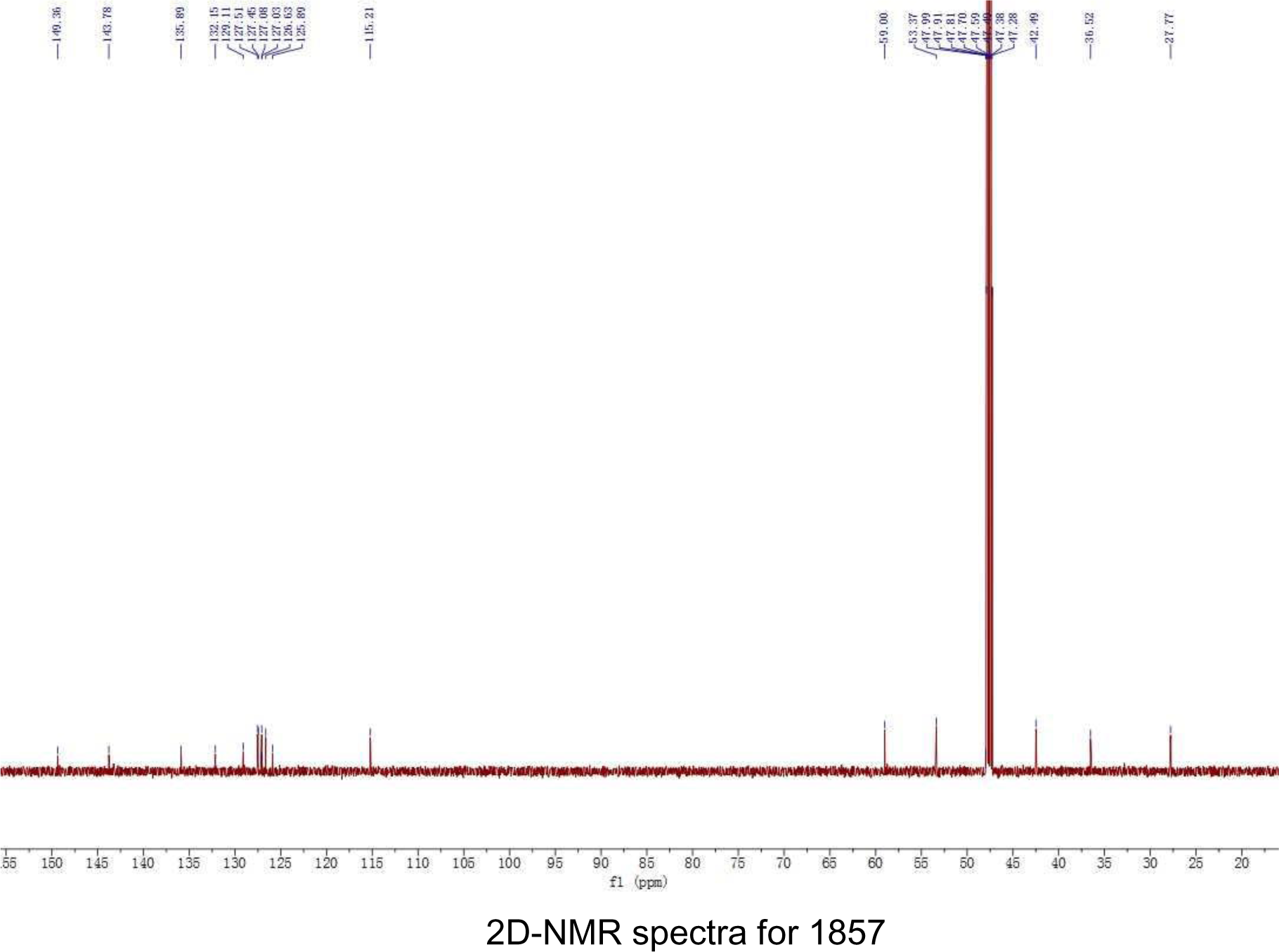
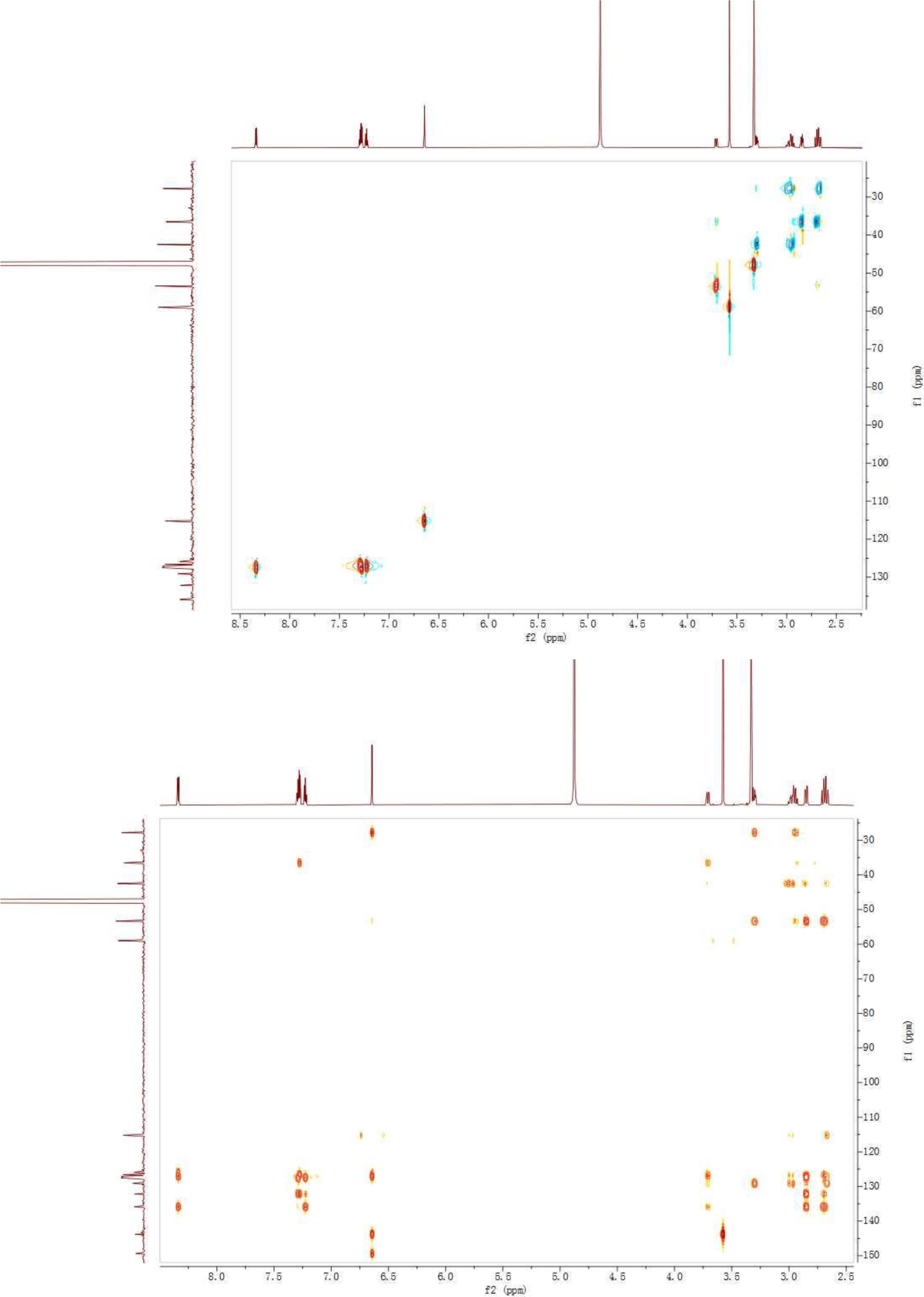
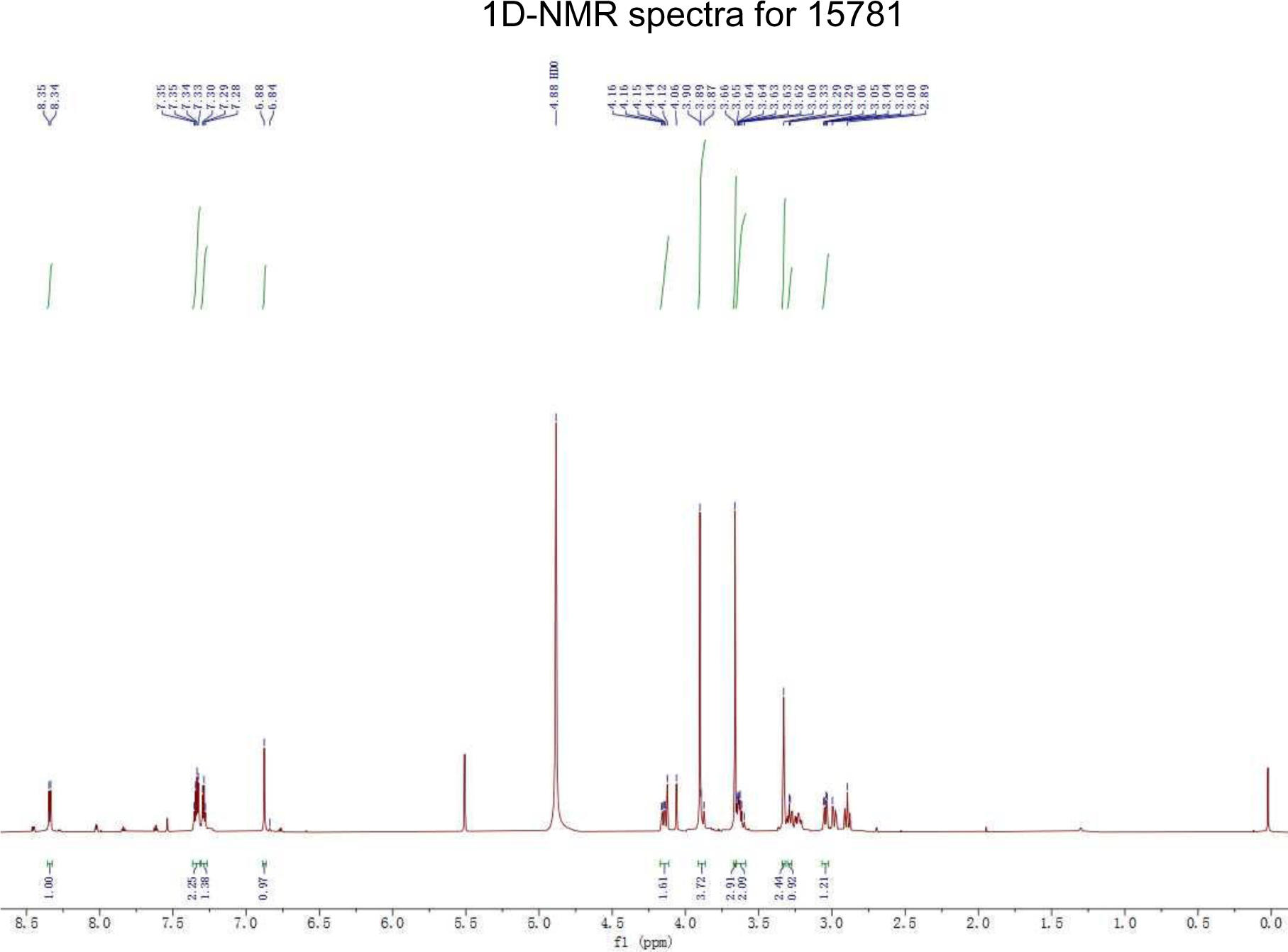
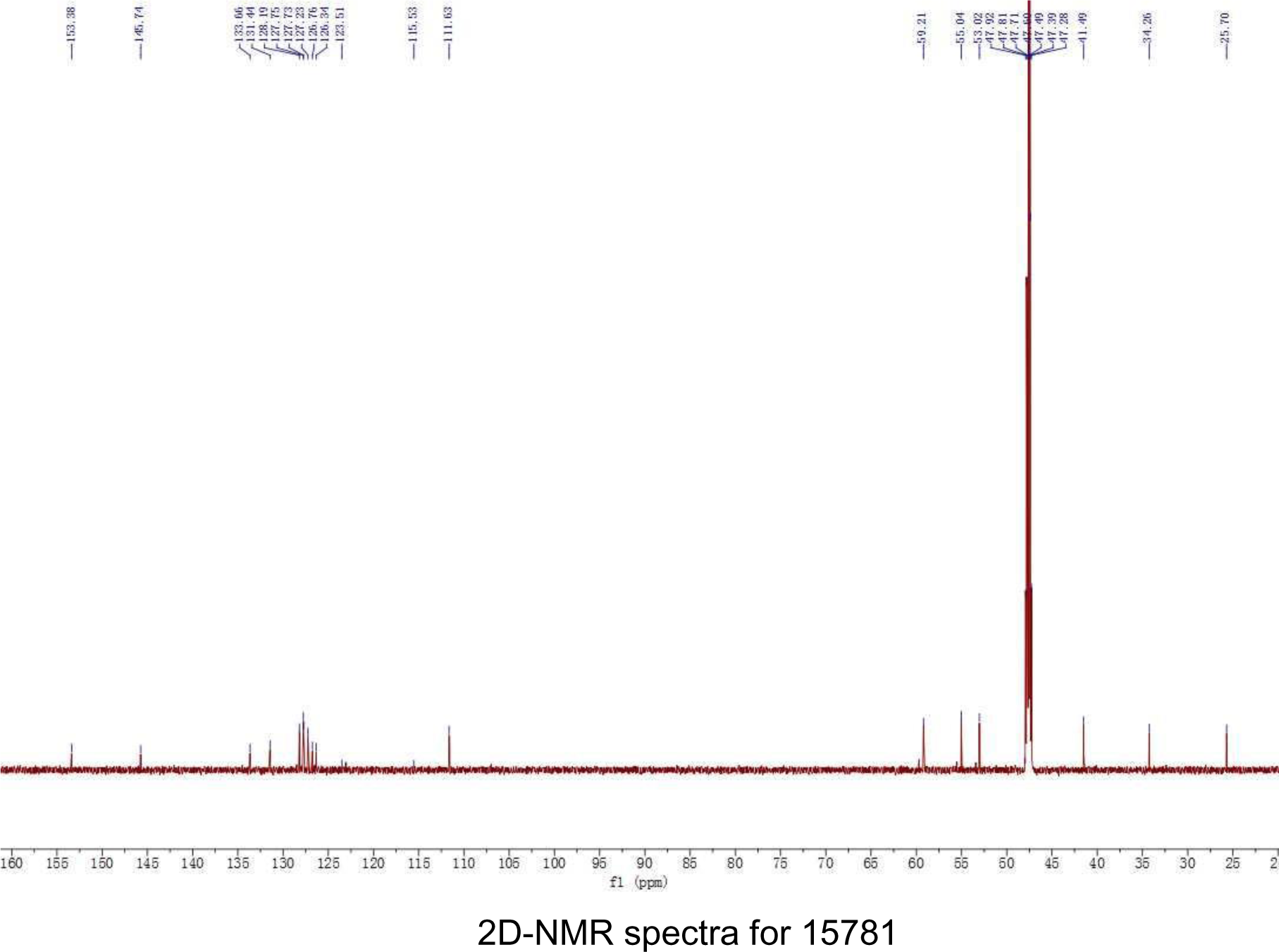
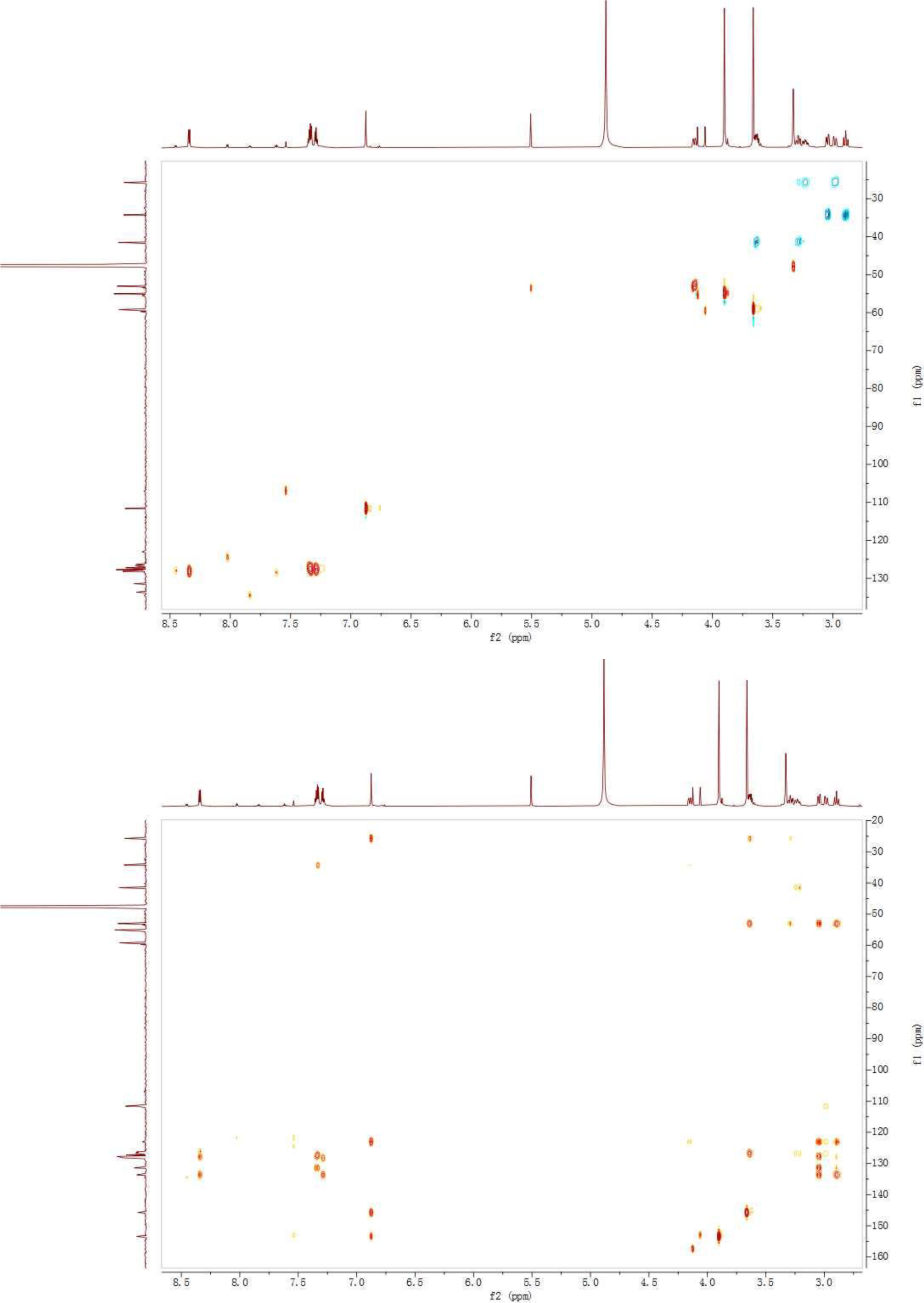

## Supplementary Note 3

### The potential role of 1857 in elucidating the structural basis of preferential G protein signaling

By virtue of the rapid advances in GPCR structural biology, the molecular basis of functional selectivity of certain biased ligands has started to unfold. In particular, Stevens and Roth groups have revealed distinct structural features of 5-HT_2B_ bound to ergotamine or LSD that are both strong β–arrestin biased agonists^10, 11^. Another elegant work on the design of β-arrestin biased ligands based on aminergic GPCR structures further pointed out that the key contacts at transmembrane helix 5 (TM5) and extracellular loop 2 (ECL2) may be responsible for G protein and β-arrestin signaling, respectively^12^. Furthermore, structural characterization of 5-HT_2B_ in complex with diverse ligands has illuminated important structural determinants essential for receptor activation and biased agonism^13^. In particular, Leu362^7.35^ in TM7 of 5-HT_2B_ appears to be a crucial determinant of preference for G protein or β-arrestin2 recruitment^13^. As for the novel biased agonist 1857 which acts on the same serotonin receptor system, we identified residues in TM5 (A222^5.46^) and TM7 (V354^7.39^) of 5-HT_2C_ that interact with the ligand and have profound effects on its biased agonism. Our finding is consistent with the previous notion that ligand engagement of TM5 and TM7 is vital to G protein biased signaling for 5-HT_2B_ and other aminergic GPCRs^12, 13^. Moreover, inability of 1857 to elicit β-arrestin2 recruitment may be attributed to its weak interaction with ECL2, a structural region posited to be critical for arrestin bias^12^. Therefore, 1857 may serve as a desirable probe for elucidating the structural basis of preferential G protein signalling and its contribution to therapeutic effects mediated by 5-HT_2C_. Of note, our docking analysis did not fully explain the regulatory roles of certain residues such as V354, which may suggest the presence of alternative conformations or dynamics of the receptor not seen in crystal structures.

---

**Supplementary Figure 1.**
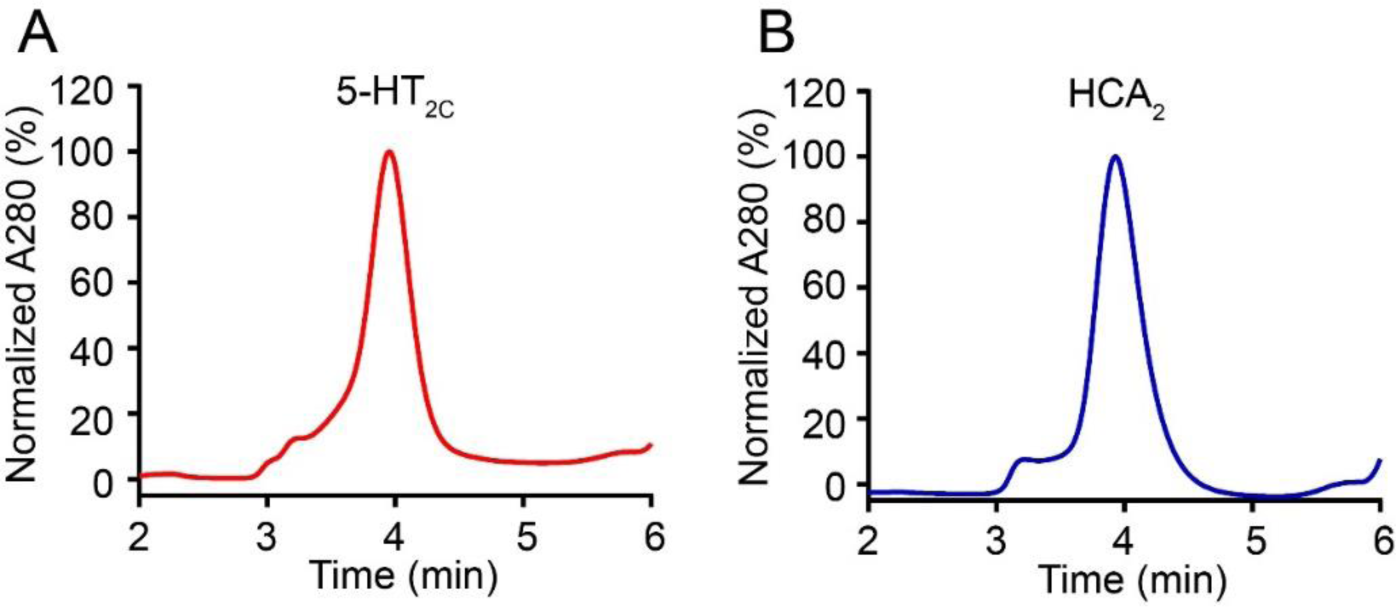
SEC analysis of purified 5-HT_2C_ and HCA2 proteins.

**Supplementary Figure 2.**
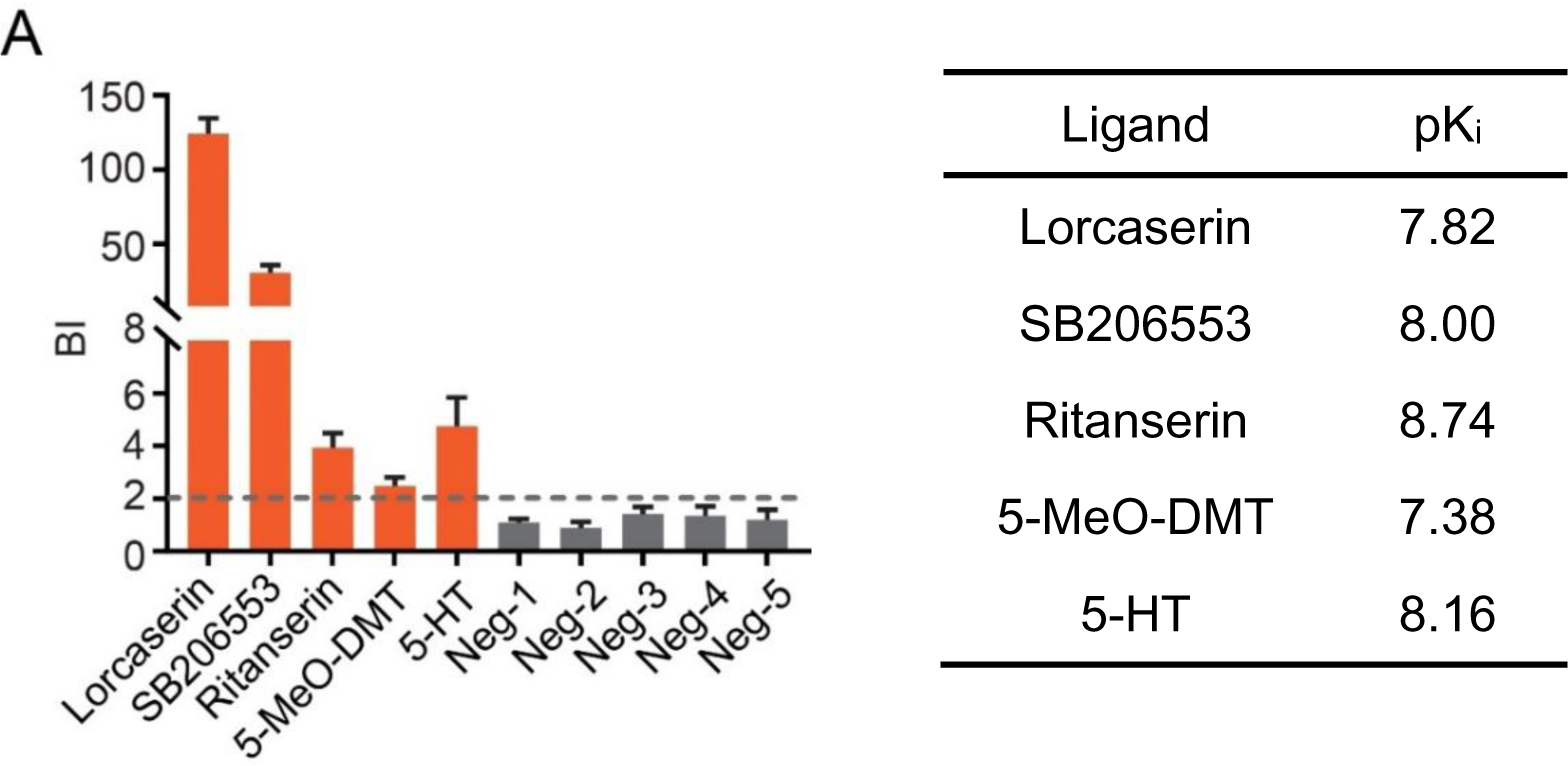
Affinity MS screening of a simple mixture of five known 5-HT_2C_ ligands (red bars) and five unrelated compounds (grey bars). Binding index (BI) refers to the ratio of MS intensity of each compound detected in the 5-HT_2C_ target *vs.* control. Data were obtained from three independent experiments. Error bars represent SEM. Binding affinity data for known ligands were obtained from ChEMBL database.

**Supplementary Figure 3.**
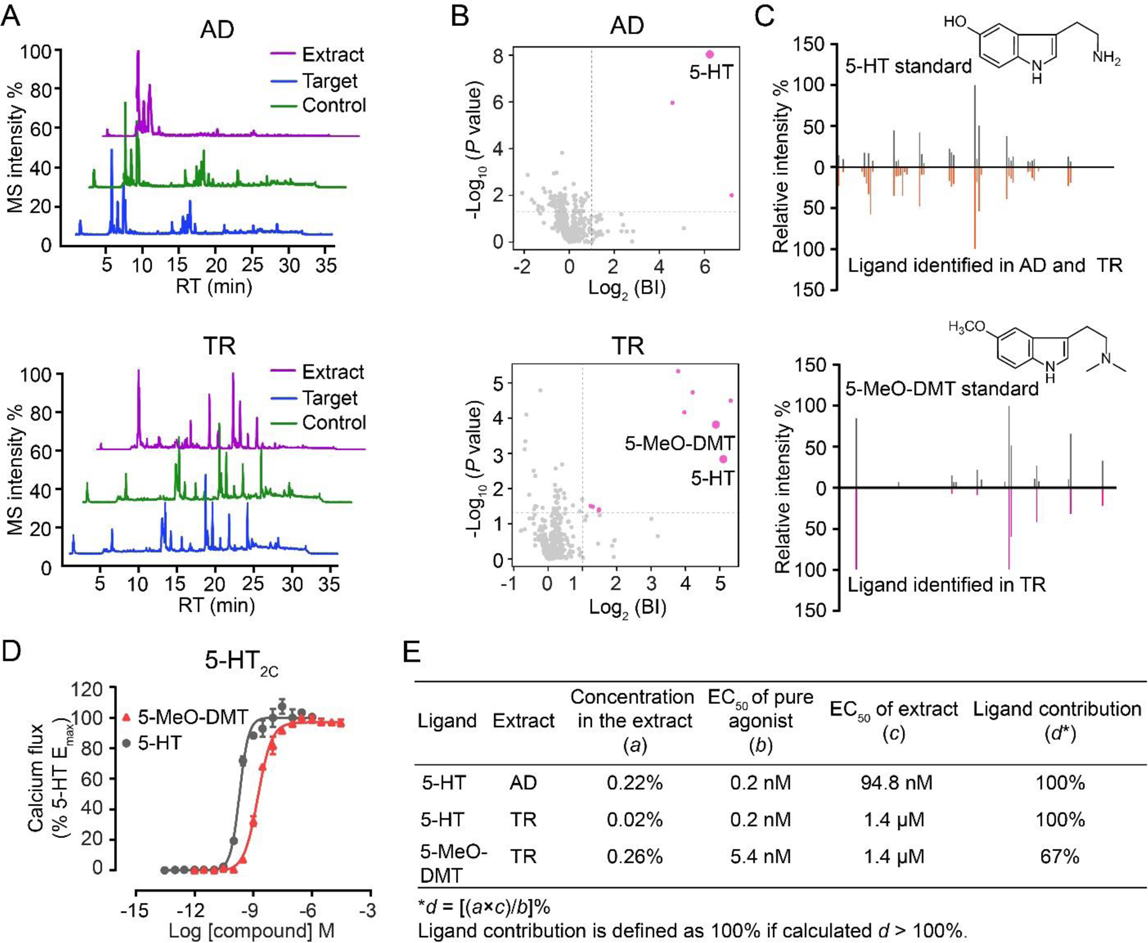
Identification of known 5-HT_2C_ agonists by screening crude extracts of *Aristolocxsshia debilis* (AD) and *Tetradium ruticarpum* (TR). (A) Representative LC-MS chromatograms of crude extract, 5-HT_2C_ target and control for AD (top) or TR (bottom). (B) Initial hits (pink dots) from screening crude extract of AD (top) or TR (bottom) by affinity MS combined with metabolomics. 5-HT and 5-MeO-DMT identified are marked by larger dots. BI, binding index. (C) Structural validation of 5-HT and 5-MeO-DMT by MSMS analysis of identified ligands and the standard. (D) 5-HT_2C_ mediated calcium mobilization elicited by 5-HT (EC_50_ = 0.2 nM) and 5-MeO-DMT (EC_50_ = 5.4 nM). (E) Calculated contribution of 5-HT and 5-MeO-DMT to the total 5-HT_2C_ activity manifested by AD and TR extracts.

**Supplementary Figure 4.**
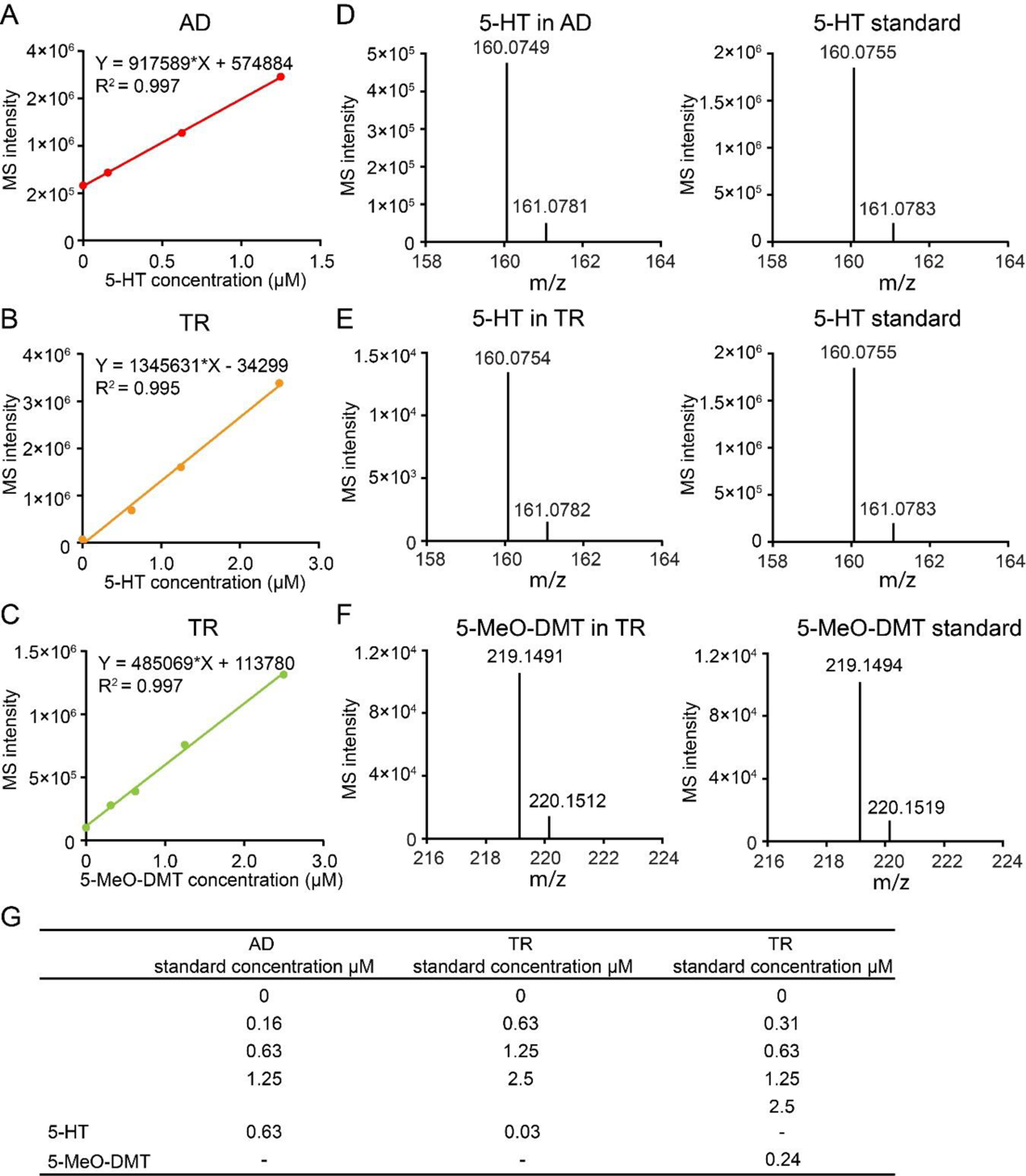
Concentrations of 5-HT and 5-MeO-DMT in the crude extracts of AD and TR determined by a standard spike-in quantification approach. MS response curves of 5-HT or 5-MeO-DMT titrated into the extracts of AD (A) or TR (B, C). Each compound was identified by HRMS measurement of the crude extract (D, E, F, left) in agreement with the standard (D, E, F, right) and retention time matching (data not shown). (G) Concentration of each compound in the original AD or TR extract was determined by extrapolating the standard curve to the origin and deriving it from the linear regression (values shown in the two rows at the bottom).

**Supplementary Figure 5.**
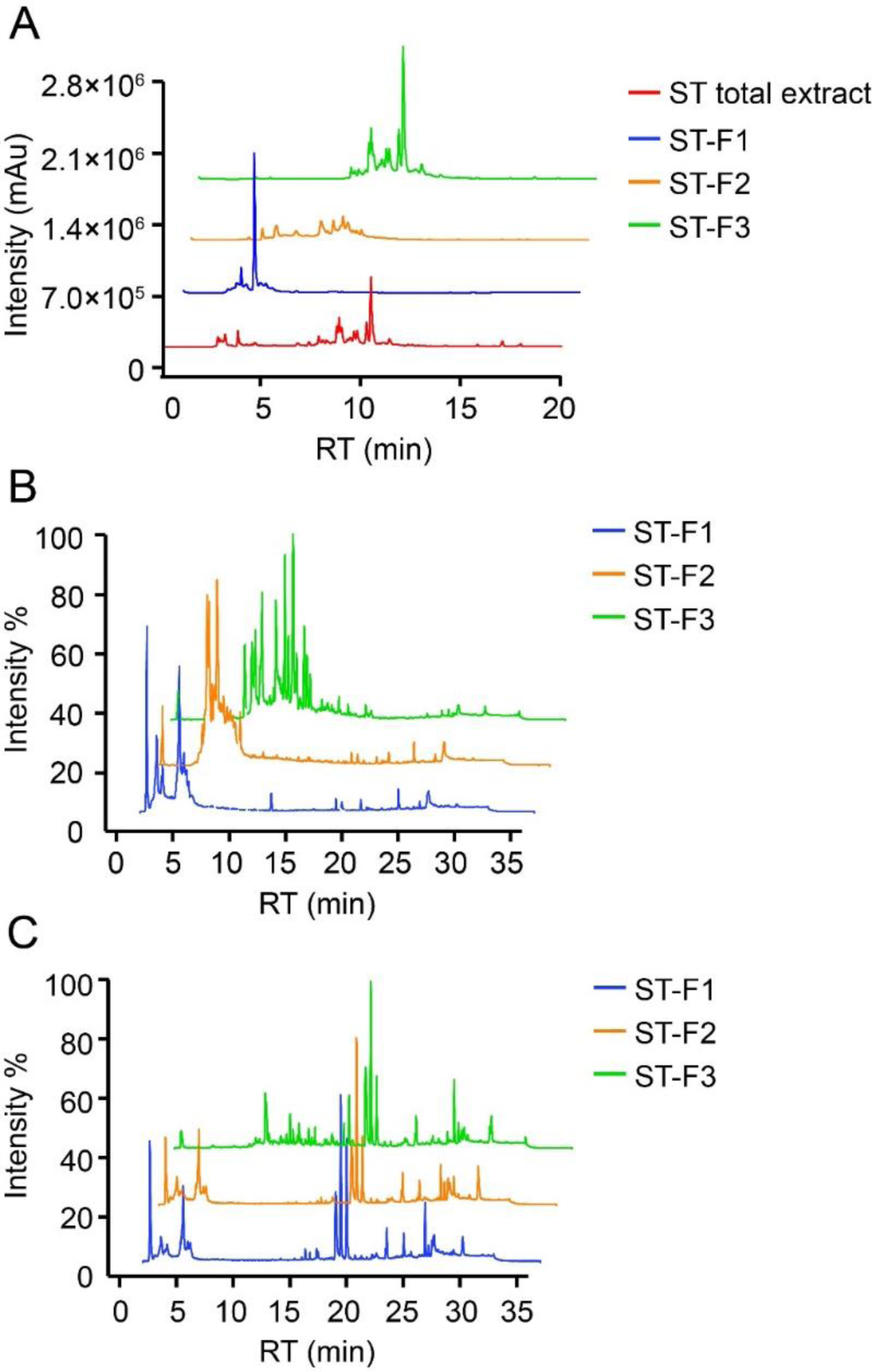
Fractionation of ST crude extract and affinity MS screening of each fraction for 5-HT_2C_ ligands. (A) LC-UV chromatograms of the total extract and its three fractions from the extract (F1, F2, F3) at 254 nm. (B) Representative LC-MS chromatograms of each ST fraction. (C) Representative LC-MS chromatograms of 5-HT_2C_ target samples from the affinity MS screening of each ST fraction.

**Supplementary Figure 6.**
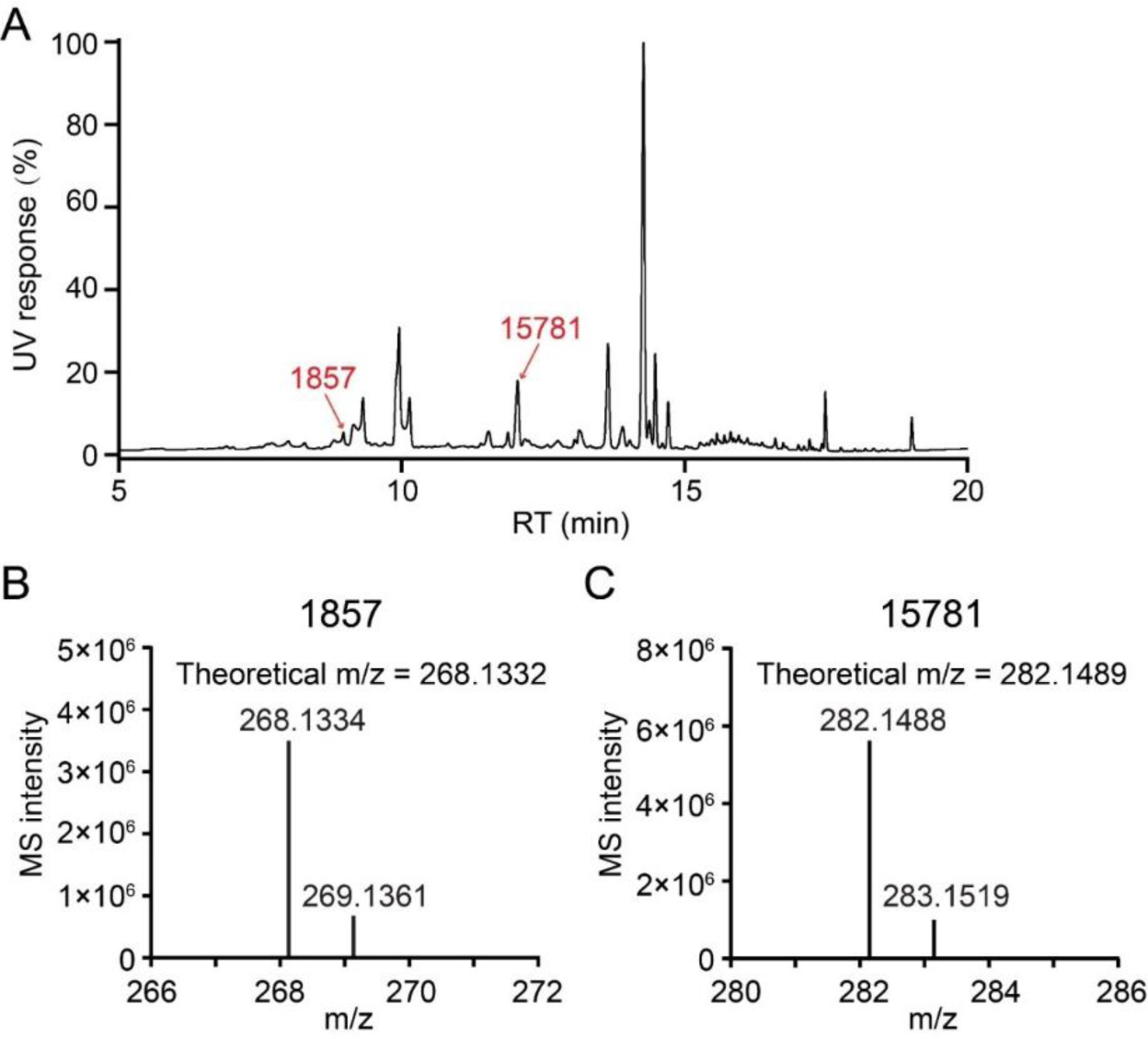
LC-UV/MS guided isolation of 1857 and 15781. (A) LC-UV chromatogram of 1857 and 15781 in the ST fraction F3 at 254 nm. (B, C) HRMS spectra of 1857 and 15781 isolated from the ST extract with measured monoisotopic mass very close to the theoretical value.

**Supplementary Figure 7.**
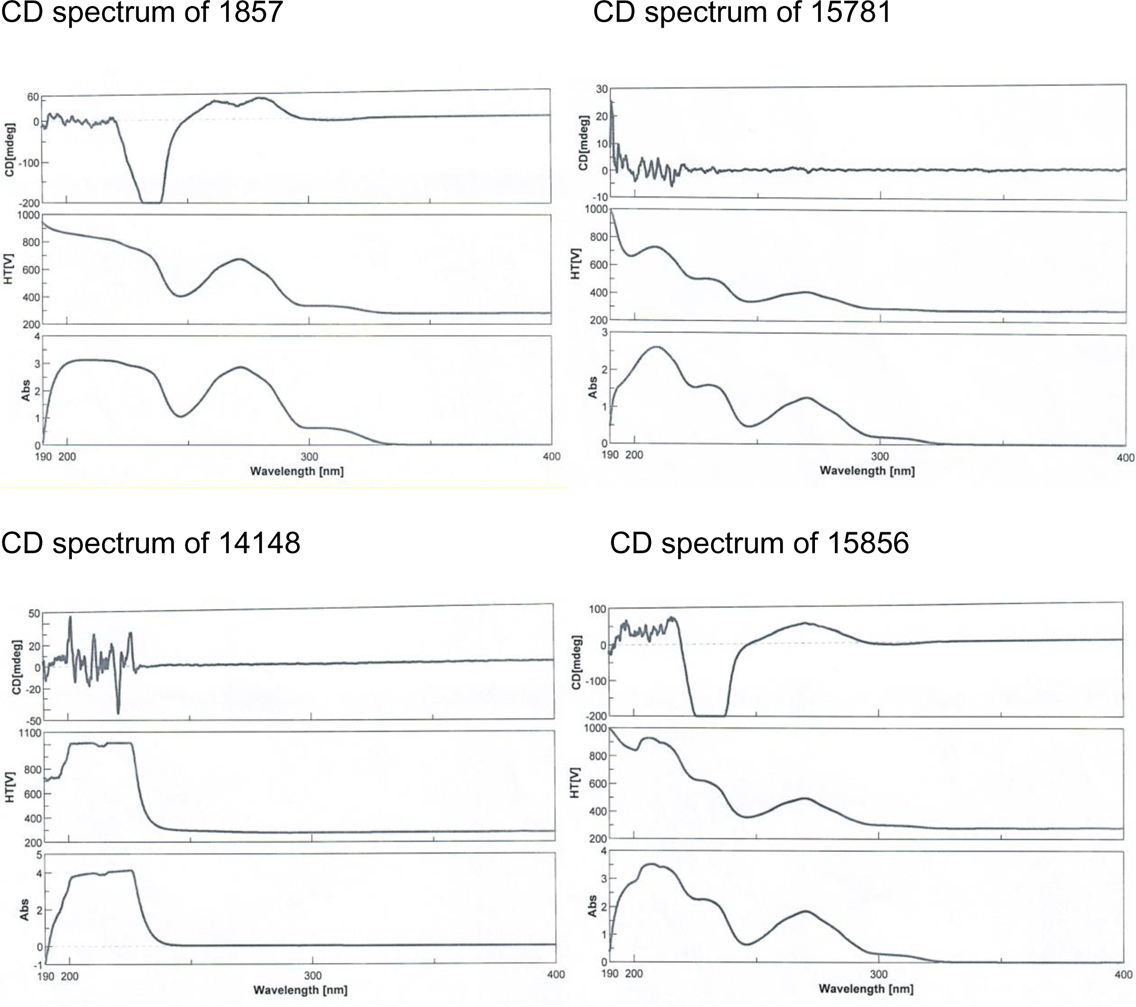
CD spectroscopy analysis of four aporphines. CD spectrum of 1857 CD spectrum of 15781

**Supplementary Figure 8.**
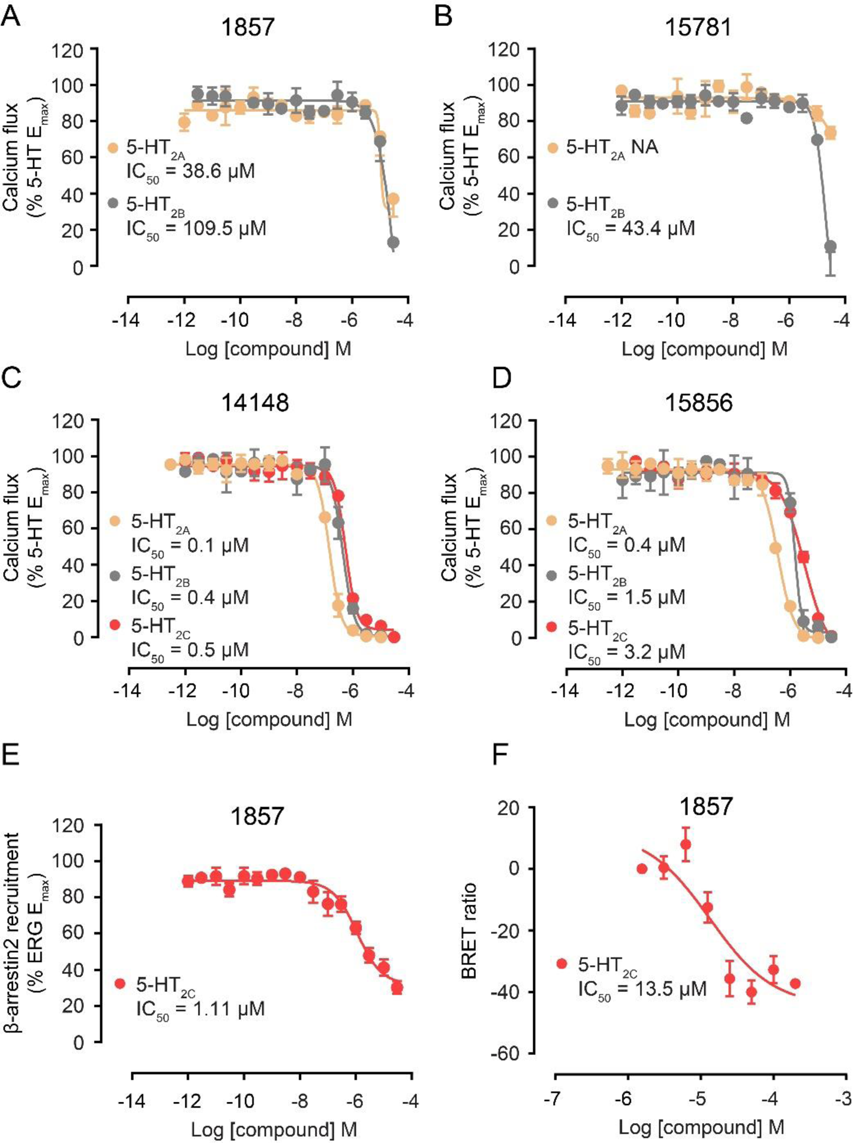
Functional profiles of four aporphines active at 5-HT_2_ subfamily members.. Weak inhibition of G_q_-mediated calcium mobilization in 5-HT_2A_ and 5-HT_2B_ expressing cells in response to 5-HT by 1857 (A) and 15781 (B). Note that 15781 did not antagonize G_q_ activity in 5-HT_2A_ expressing cells under the same condition. Non-selective inhibition of G_q_-mediated calcium mobilization in 5-HT_2A_, 5-HT_2B_ and 5-HT_2C_ expressing cells in response to 5-HT by 14148 (C) and 15856 (D). Inhibition of β-arrestin2 recruitment in 5-HT_2C_ expressing cells by 1857 as measured by Tango (E) or BRET (F) assays. Data represent means ± SEM of three independent experiments performed in triplicate. NA, no measurable activity.

**Supplementary Figure 9.**
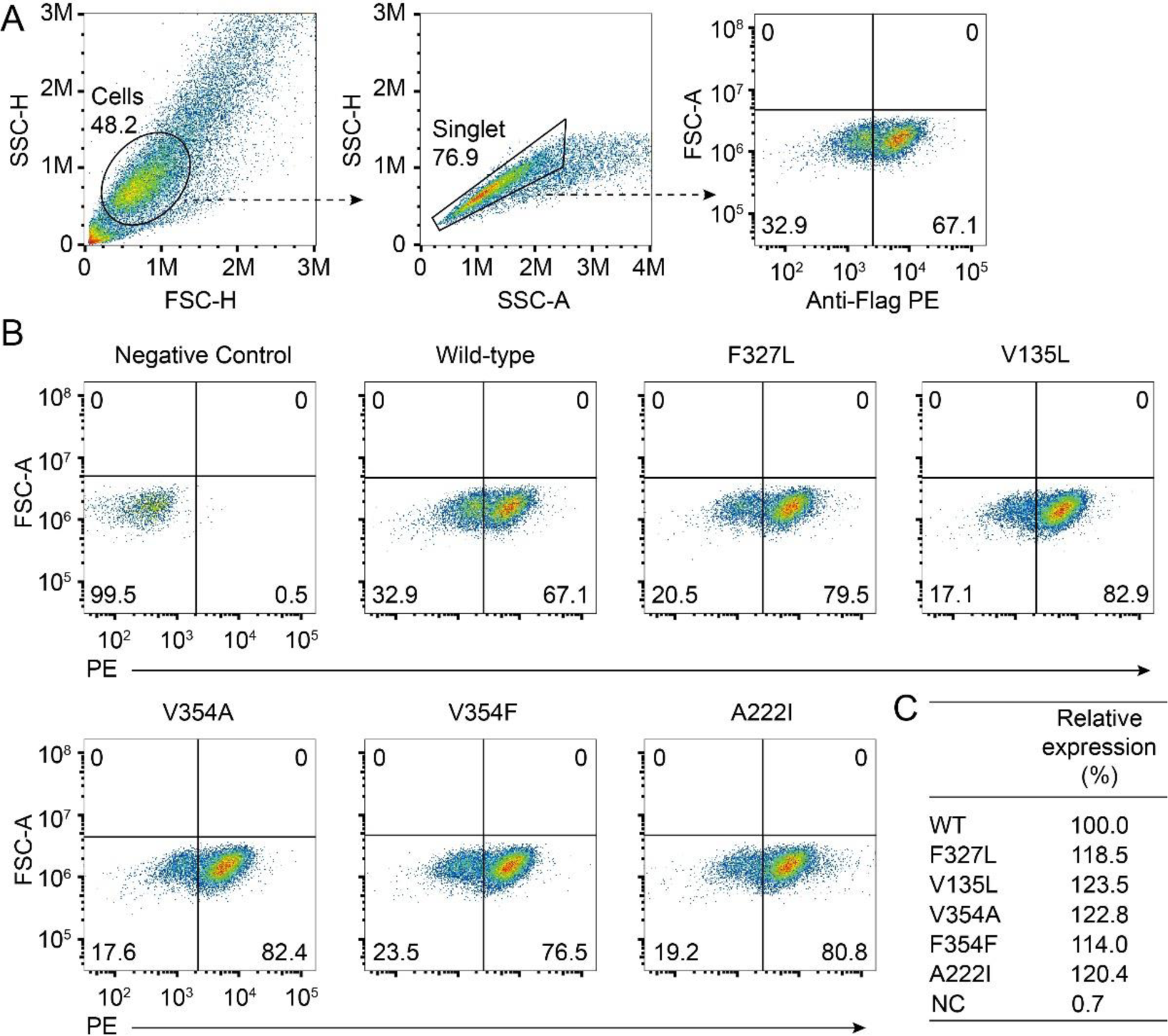
Cell surface expression of wild-type and mutant 5-HT_2C_ receptors. (A) Gating strategy of cell sorting. (B) FACS analysis of wild-type and mutant 5-HT_2C_ receptors. (C) Relative surface expression of wild-type and mutant 5-HT_2C_ receptors.

**Supplementary Table 3.**
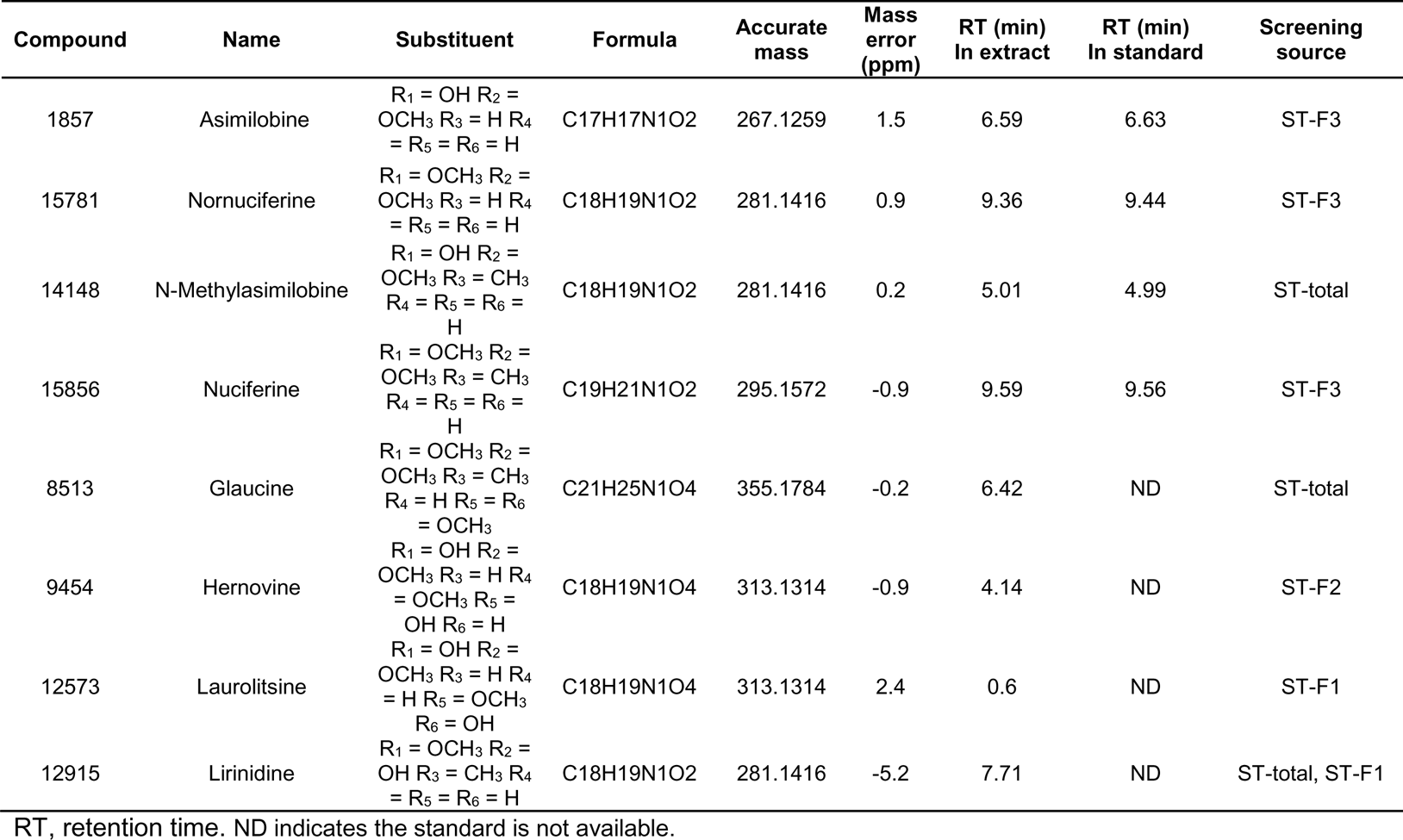
Identification of eight aporphine analogues from Stephania tetrandra (ST) as putative 5-HT_2C_ ligands. Each compound was identified from screening the total extract (ST-total) or a specific fraction of the extract (ST-F1/F2/F3) as indicated in “Screening source”.

**Supplementary Table 4.** K_i_ values of four aporphines and lorcaserin at 5-HT_2A_/2B/2C receptors.

| Compound | h5-HT<sub>2A</sub> [<sup>3</sup>H]-Ketanserin<br>K<sub>i</sub> nM (pK<sub>i</sub> ± SEM) | h5-HT<sub>2B</sub> [<sup>3</sup>H]-LSD<br>K<sub>i</sub> nM (pK<sub>i</sub> ± SEM) | h5-HT<sub>2C</sub> [<sup>3</sup>H]-Mesulergine<br>K<sub>i</sub> nM (pK<sub>i</sub> ± SEM) |
| --- | --- | --- | --- |
| 1857 | 375.9 (6.43 ± 0.08) | 74.2 (7.13 ± 0.05) | 193.9 (6.71 ± 0.04) |
| 15781 | 925.0 (6.03 ± 0.06) | 263.5 (6.58 ± 0.06) | 446.2 (6.35 ± 0.05) |
| 14148 | 60.0 (7.22 ± 0.04) | 3.38 (8.47 ± 0.03) | 20.9 (7.68 ± 0.03) |
| 15856 | 149.7 (6.83 ± 0.05) | 14.3 (7.85 ± 0.03) | 22.2 (7.65 ± 0.04) |
| Lorcaserin | 765.1 (6.11 ± 0.06) | 140.2 (6.85 ± 0.05) | 123.1 (6.91 ± 0.05) |

Data represent K<sub>i</sub> and pK<sub>i</sub> from three independent experiments.

**Supplementary Table 5.**
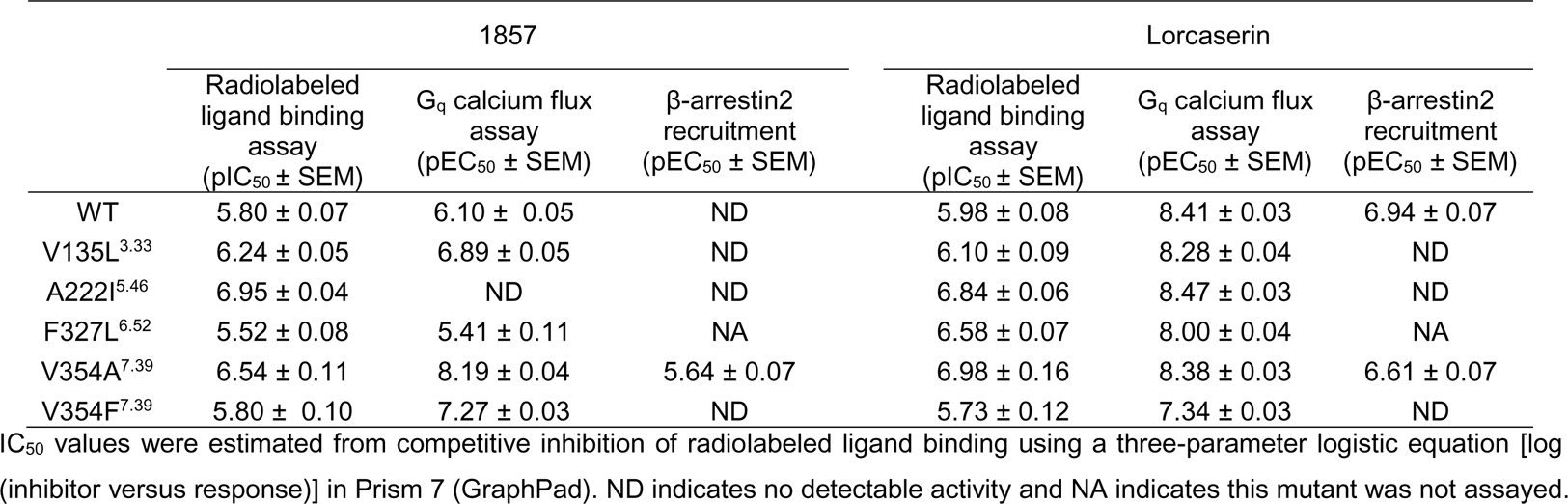
IC50/EC_50_ values of 1857 and lorcaserin at wild-type and mutant 5-HT_2C_ receptors.

|  | 1857 |  |  | Lorcaserin |  |  |
| --- | --- | --- | --- | --- | --- | --- |
|  | Radiolabeled<br>ligand binding<br>assay<br>(pIC<sub>50</sub> ± SEM) | G<sub>q</sub> calcium flux<br>assay<br>(pEC<sub>50</sub> ± SEM) | β-arrestin2<br>recruitment<br>(pEC<sub>50</sub> ± SEM) | Radiolabeled<br>ligand binding<br>assay<br>(pIC<sub>50</sub> ± SEM) | G<sub>q</sub> calcium flux<br>assay<br>(pEC<sub>50</sub> ± SEM) | β-arrestin2<br>recruitment<br>(pEC<sub>50</sub> ± SEM) |
| WT | 5.80 ± 0.07 | 6.10 ± 0.05 | ND | 5.98 ± 0.08 | 8.41 ± 0.03 | 6.94 ± 0.07 |
| V135L<sup>3.33</sup> | 6.24 ± 0.05 | 6.89 ± 0.05 | ND | 6.10 ± 0.09 | 8.28 ± 0.04 | ND |
| A222I<sup>5.46</sup> | 6.95 ± 0.04 | ND | ND | 6.84 ± 0.06 | 8.47 ± 0.03 | ND |
| F327L<sup>6.52</sup> | 5.52 ± 0.08 | 5.41 ± 0.11 | NA | 6.58 ± 0.07 | 8.00 ± 0.04 | NA |
| V354A<sup>7.39</sup> | 6.54 ± 0.11 | 8.19 ± 0.04 | 5.64 ± 0.07 | 6.98 ± 0.16 | 8.38 ± 0.03 | 6.61 ± 0.07 |
| V354F<sup>7.39</sup> | 5.80 ± 0.10 | 7.27 ± 0.03 | ND | 5.73 ± 0.12 | 7.34 ± 0.03 | ND |

IC<sub>50</sub> values were estimated from competitive inhibition of radiolabeled ligand binding using a three-parameter logistic equation [log (inhibitor versus response)] in Prism 7 (GraphPad). ND indicates no detectable activity and NA indicates this mutant was not assayed.

**Supplementary Table 6.** Blood-brain barrier penetrance of 1857 in mice.

| Sample | Plasma | Brain | Ratio<br>($C_{\text{brain}}/C_{\text{plasma}}$) |
| --- | --- | --- | --- |
|  | Conc. (ng/mL) | Conc. (ng/g) |  |
| 1857-30 min | 659.7 ± 176.7 | 11060.3 ± 2774.9 | 16.8 |
| 1857-240 min | 36.7 ± 3.4 | 236.0 ± 33.1 | 6.4 |

